# The crystal structure of the varicella zoster Orf24-Orf27 nuclear egress complex spotlights multiple determinants of herpesvirus subfamily specificity

**DOI:** 10.1101/2021.08.23.457313

**Authors:** Johannes Schweininger, Mark Kriegel, Sigrun Häge, Marcus Conrad, Sewar Alkhashrom, Josephine Lösing, Sigrid Weiler, Julia Tillmanns, Claudia Egerer-Sieber, Andrea Decker, Tihana Lenac Roviš, Jutta Eichler, Heinrich Sticht, Manfred Marschall, Yves A. Muller

**Affiliations:** Division of Biotechnology, Department of Biology, Friedrich-Alexander-Universität Erlangen-Nürnberg (FAU), Erlangen, Germany; Institute for Clinical and Molecular Virology, Friedrich-Alexander-Universität Erlangen-Nürnberg (FAU), Medical Center Erlangen, Germany; Division of Bioinformatics, Institute of Biochemistry, Friedrich-Alexander-Universität Erlangen-Nürnberg (FAU), Erlangen, Germany; Department of Chemistry and Pharmacy, Friedrich-Alexander-Universität Erlangen-Nürnberg (FAU), Erlangen, Germany; Center for Proteomics, Faculty of Medicine, University of Rijeka, Rijeka, Croatia

## Abstract

Varicella zoster virus (VZV) is a human pathogen from the α-subfamily of herpesviruses. Here, the crystal structure of the VZV Orf24-Orf27 complex is described, representing the essential viral core nuclear egress complex (NEC) that orchestrates the egress of the preassembled capsids from the nucleus. While previous studies have primarily emphasized the finding that the architecture of core NEC complexes is highly conserved among herpesviruses, the present report focusses on subfamily-specific structural and functional features that help explain the differences in the autologous *versus* nonautologous interaction patterns observed for NEC formation across herpesviruses. CoIP and confocal imaging data show that Orf24-Orf27 complex formation displays some promiscuity in a herpesvirus subfamily-restricted manner. At the same time, analysis of the NEC formation thermodynamic parameters of three prototypical α-, β- and γ herpesviruses, i.e. VZV, human cytomegalovirus (HCMV) and Epstein-Barr virus (EBV) reveals highly similar binding affinities for the autologous interaction with some specific differences in the enthalpy and entropy terms. Computational alanine scanning and structural comparisons highlight intermolecular interactions shared among α-herpesviruses that are clearly distinct from those seen in β- and γ-herpesviruses. Combined, these data allow to explain the distinct properties of specificity and permissivity so far observed in herpesviral NEC interactions. These findings might prove highly valuable when attempting to target multiple herpesvirus core NECs with selective or broad-acting drug candidates.

## INTRODUCTION

Members of the family *Herpesviridae* are membrane-enveloped viruses, characterized by a comparably large particle size (100-300 nm) and a linear double-stranded DNA genome. The nine presently known human herpesviruses can be assigned to three subfamilies, i.e. the α-, β- and γ-*Herpesvirinae*, based on their morphology, genetics and biological properties (1). A typical feature of all herpesvirus infections is their lifelong persistence in the host with extended periods of latency followed by intermittent periods of reactivation. While most herpesviral infections may remain on subclinical or asymptomatic stages in immunocompetent persons, severe impairments of health can occur in older persons and/or those with a compromised immune system, such as patients suffering from AIDS/immunodeficiency virus type 1 infection (HIV-1), immunosuppressive treatment or cancer therapy. The present study focusses on the α-herpesvirus varicella zoster virus (VZV, HHV-3), which shows a seroprevalence of >90 % in the human population worldwide (2). VZV causes chickenpox (*varicella*) in children and persists in the nervous system of the immunocompetent host during latency. Upon reactivation, VZV leads to lesions known as shingles (*zoster*) and can cause severe neurological conditions such as acute sequelae or continuing burning pain. The prototype viruses of the β- and γ-subfamilies are human cytomegalovirus (HCMV) and Epstein-Barr virus (EBV), respectively. HCMV has a seroprevalence of 40 % to 95 % in various parts of the world and congenital HCMV infections (cCMV) acquired during pregnancy can lead to stillbirth, severe birth defects in newborns as well as late-onset developmental retardation (3–5). EBV is the causative agent of infectious mononucleosis, and in addition EBV infections are associated with various, typically malignant tumors, such as Burkitt lymphoma, Hodgkin’s lymphoma, post-transplant B and T cell lymphoma, nasopharyngeal and gastric carcinoma (6). Animal herpesviruses such as bovine α-herpesvirus 1, equine herpesviruses or pseudorabies virus (PRV) can cause pathogenic infections in domesticated animals. These are often met by a strict culling policy causing severe economic repercussions (7–9). Although antiviral drugs and vaccines exist for combatting some herpesvirus infections, approved vaccines against HCMV and EBV are still missing, and in cases of clinically problematic VZV infections, antiviral therapy has not yet achieved the desired efficacy. Treatments are also under the continuous threat of the emergence of viral drug resistance or vaccine escape mutations. Hence, the identification of novel antiherpesviral drugs and targeting strategies remains a global research focus (10, 11).

Common to all herpesviruses is the preassembly of capsids in the nucleus, whereas the final maturation of virions occurs in the cytosol and trans-Golgi membrane vesicles. Due to size constraints, intranuclear capsids cannot migrate from the nucleoplasm to the cytosol *via* the nuclear pore complex (NPC) but rely on the mechanism of nuclear egress that is regulated in a herpesvirus-specific manner (12, 13). This process starts with the formation of the core nuclear egress complex (core NEC) formed between two viral proteins, namely between a membrane-associated and a nucleoplasmic-soluble protein. The core NEC combines multiple regulatory aspects. For once, it functions as a scaffold platform for the recruitment of virus- and host cell-derived NEC-associated effector proteins that destabilize the lamina at the inner layer of the nuclear envelope and induce membrane fission (13, 14). Secondly, the complex associates to higher oligomers, specifically to hexameric and possibly pentameric arrangements, that induce capsid docking followed by budding through the inner nuclear membrane (15). Experimental structural data on the core NECs are so far available from the α-herpesviruses herpes simplex virus type 1 (HSV-1) and PRV, the β-herpesvirus HCMV and the γ-herpesvirus EBV (Table S1) (16–20). A hallmark of all complexes is the so-called hook-into-groove interaction with the membrane-associated NEC protein (e.g. VZV Orf24) forming a groove-like interaction surface, onto which a contiguous segment from the nucleoplasmic-soluble NEC protein (e.g. VZV Orf27) binds in a hook-like manner (3,13,21). The hook-like segment is approximately thirty residues long, is formed by two α-helices and interacts with high affinity with the groove protein thus resulting in core NEC assembly (19). In all known characterized complexes, this interaction motif accounts for about 80 % of the total number of interactions formed between the hook and the groove protein. The hook-into-groove interaction principle of NEC formation appears highly specific for individual herpesviruses. Interestingly, however, an experimental testing of selected examples of NEC protein interactions between herpesviruses in a nonautologous, cross-viral manner, revealed some promiscuity of interaction as observed for NEC protein pairs derived from members of the same subfamily (22). Because of the scarcity of currently available structural information, the structural determinants for the subfamily-specific interaction pattern remains currently purely understood.

**Table 1.** Binding parameters between hook and groove proteins in VZV, HCMV and EBV.

|  | ITC measurements |  |  |  |  |  | ELISA |
| --- | --- | --- | --- | --- | --- | --- | --- |
| | Protein titrated into the cell | Protein in cell | $K_d$ [nM]<sup>a</sup> | n | $\Delta H$ [kJ/mol] | $-T\Delta S$ [kJ/mol]<sup>b</sup> | $EC_{50}$ [nM] |
| VZV | Orf24 | Orf27 | $11.5 \pm 0.2$ | 0.6 | $-91.9 \pm 1.0$ | $46.6 \pm 1.0$ | 49.2 |
| HCMV | pUL50 | pUL53 | $19.8 \pm 4.6$ | 0.9 | $-27.0 \pm 0.1$ | $-17.0 \pm 0.5$ | 11.6 |
| EBV | BFRF1 | BFLF2 | $85.6 \pm 40.5$ | 0.7 | $-63.3 \pm 0.5$ | $22.8 \pm 1.5$ | 34.4 |

<sup>a</sup> Experiments were performed in triplicates.
<sup>b</sup> $T = 298.15$ K

Here, we report the 2.1 Å resolution crystal structure of the core NEC complex from α-herpesvirus VZV, namely the complex formed between the grove protein Orf24 and the hook protein Orf27. Computational alanine scanning and structural comparisons between VZV and the previously reported α-herpesviral NECs of HSV-1 and PRV revealed a number of shared intermolecular interaction features that clearly differed from those in the β-/γ-herpesviral NECs of HCMV and EBV, respectively. Referring to functional aspects, an analysis by coimmunoprecipitation (CoIP) and confocal imaging (CLSM) colocalization showed that VZV Orf24-Orf27 complex formation displays some promiscuity regarding binding to other α-herpesviral NEC protein binding partners, while this is not the case with regard to nonautologous interactions between core NEC proteins belonging to different herpesviral subfamilies. These findings thus confirm initial data reported earlier and explain both, the distinct properties and specificity restrictions so far observed by various studies on herpesviral NEC interactions.

## RESULTS

### Molecular characteristics of the α-, β- and γ-herpesviral core NEC proteins as expressed in cell culture-based viral infection models

We used α- (VZV), β- (HCMV) and γ-herpesviruses (EBV) that express the GFP (green fluorescent protein) as a reporter for monitoring the courses of an *in vitro* infection, in order to determine the expression characteristics of core nuclear egress proteins. Infected cells were used for collecting consecutive samples at 2, 4 and 6 days post-infection (d p.i.) to analyze viral replication kinetics in a comparative manner. Stocks of VZV (cell-associated virus inoculi) and HCMV (cell-free supernatant virus inoculi) were used for fresh infection of primary human foreskin fibroblasts (HFFs) and EBV-positive producer cells (Akata-BX1 carrying EBV-GFP) were induced for the viral lytic cycle by chemical stimulation with TPA. Samples were analyzed by Western blot (Wb) staining procedures using a panel of virus-specific monoclonal antibodies, i.e. those directed to the viral core NEC proteins as well as the respective egress-regulating viral protein kinases (in addition to control proteins of infection such as the GFP reporter and HCMV-/EBV-encoded immediate early proteins IE1p72 or BZLF1, respectively). The NEC proteins and NEC-relevant kinases of all three viruses were expressed with early-late kinetics, mostly starting with detectable signals at 2 d p.i. in dependence on the multiplicity of infection (MOI) or the concentration of TPA induction (Fig. S1). The detected NEC proteins comprised molecular masses between 30-42 kDa (VZV, Orf24 30 kDa and Orf27 27-34 kDa; HCMV, pUL50 42 kDa and pUL53 38 kDa; EBV, BFRF1 37-38 kDa), while the protein kinases ranged to higher molecular-mass sizes between 42-100 kDa (the latter showing the bands of three isoforms of HCMV pUL97 70-100 kDa, (23)). Notably, several of these viral proteins were expressed in more than one distinct variety, thus also including additional bands derived from protein phosphorylation and other posttranslational modifications (Fig. S1) (13,23–28). The NEC association of regulatory viral protein kinases provides an essential effector function for all so far analyzed α-, β- and γ-herpesviral multicomponent NECs, as previously reported by our group and others (13,26,29).

### The VZV Orf24-Orf27 heterodimeric complex is formed from monomeric protomers and displays nM affinity

Individually purified Orf24 and Orf27 proteins from VZV behave as monomers in solution and readily form a stable heterodimeric complex upon mixing (Fig. 1). This behavior, namely monomeric protomers yielding a heterodimeric core NEC, has already been reported for the α-herpesvirus core NEC formed by the HSV-1 proteins pUL34 (groove protein) and pUL31 (hook protein) (30). However, in contrast to α-herpesviral pUL31 proteins from HSV-1 and PRV, which cannot be studied in solution in absence of a solubility tag, the pUL31-homologous protein of VZV, i.e. Orf27, behaves differently (16, 20). Both Orf24 and Orf27 remain soluble after removal of the His- and GST-tag, respectively, and display circular dichroism spectra of well-folded proteins (data not shown).

**Figure 1.**
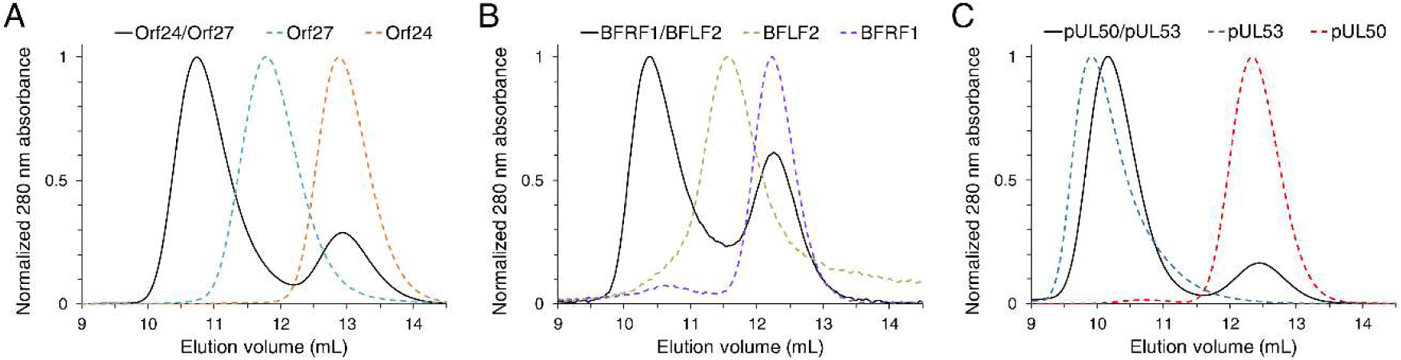
Stoichiometry of core NEC formation. A, Core NEC formation in VZV in comparison to complex formation in B, EBV and C, HCMV as analyzed using a gel filtration experiment. A, VZV Orf24 (residues 16-189, orange, dashed line), Orf27 (77-333, cyan, dashed line) or both (black, solid line). B, EBV BFRF1 (1-192, purple, dashed line), BFRF2 (78-318, green, dashed line) or both (black, solid line). C, HCMV pUL50 (1-175, red, dashed line), pUL53 (50-292, blue, dashed line) or both (black, solid line).

To shed light on possible species-related features as well as subfamily-specific differences in core NEC formation, we purified the individual proteins from VZV, HCMV and EBV and investigated complex formation using identical experimental setups. Size exclusion chromatography experiments show that in addition to VZV, the EBV BFRF1-BFLF2 complex formation also originates from the interaction of monomeric proteins. In contrast, the HCMV pUL50-pUL53 heterodimeric complex is formed upon recombination of monomeric pUL50 with homodimeric pUL53 (Fig. 1, Fig. S2) (31). As previously reported for HCMV, only the groove protein pUL50 behaves as a monomer in solution, whereas the hook protein pUL53 is dimeric, at least in vitro with bacterially-produced truncated pUL53 (31). Moreover, the hook segment of pUL53 is responsible for pUL53 homodimerization (17, 31). Interestingly however, homodimerization could not be observed under conditions of eukaryotic expression and the use of various interaction studies (32).

To gain insight into the thermodynamic profiles of the interactions of the three prototypical subfamily members, the affinities of the VZV, EBV and HCMV core NEC protein complexes were measured by isothermal titration calorimetry (ITC) and ELISA experiments. In both assay formats, affinities in the mid-nanomolar range (10 – 90 nM) were determined for all three interactions, indicating similar stabilities of the complexes. (Table 1, Fig. 2, Fig. S3).

**Figure 2.**
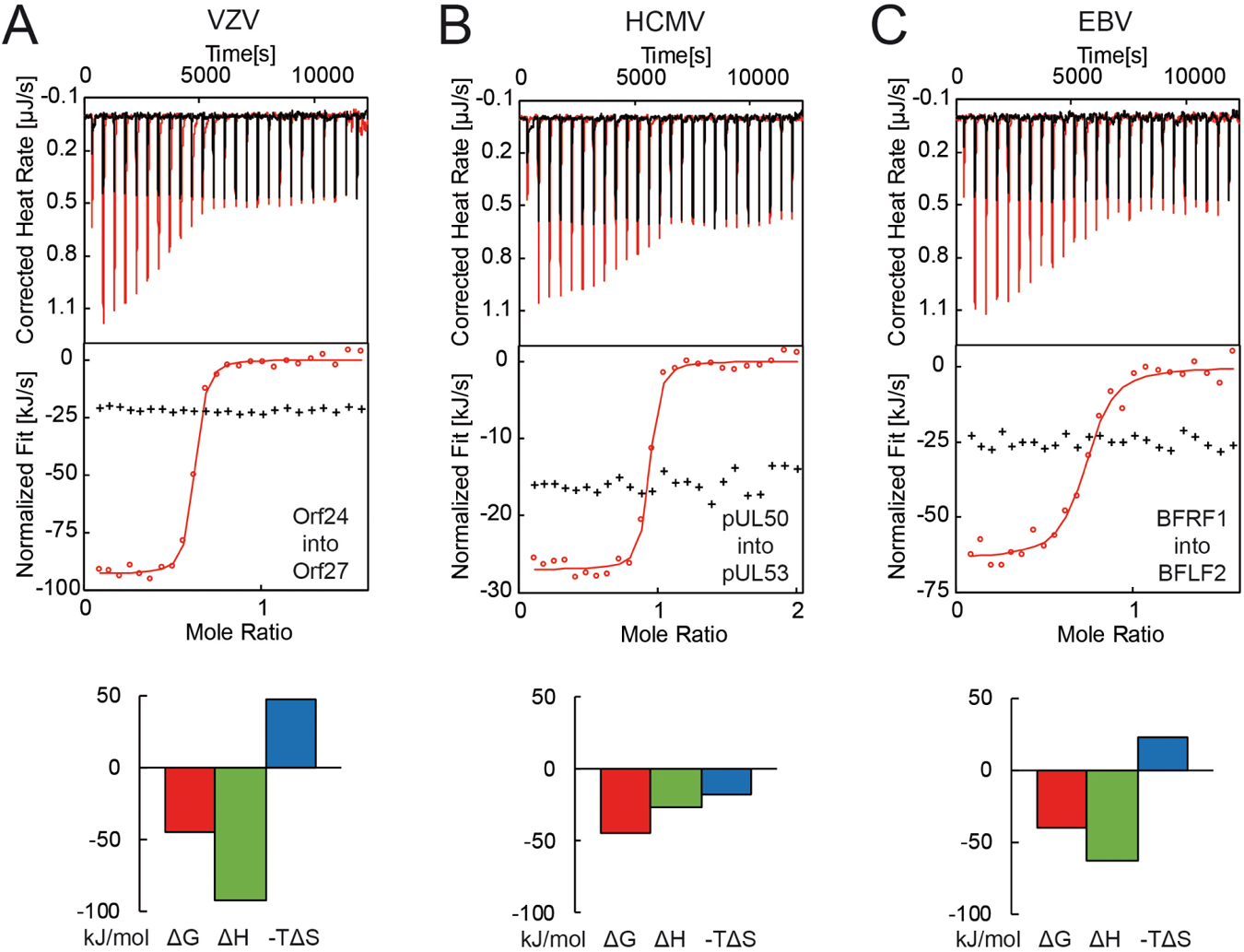
Determination of the thermodynamic parameters of VZV ORF24-ORF27 complex formation in comparison to HCMV and EBV via ITC. A, VZV Orf24-Orf27, B, HCMV pUL50-pUL53 and C, EBV BFRF1-BFLF2 complex formation. ITC traces, the integrated heats (circles) and the fitted binding models (lines) of the NEC formations are shown in red, the corresponding blank titrations in black (crosses).

The EBV and HCMV affinities are slightly higher than those previously observed for the interaction of HCMV pUL50 and EBV BFRF1 with peptides presenting the hook segments of HCMV pUL53 and EBV BFLF2, respectively. The peptides only bound with 120 nM (EBV) and 117 nM (HCMV) affinity (3). This can be easily explained by the additional contacts observed and anticipated in the complexes of the entire proteins, namely contacts formed in addition to the hook-into-groove interaction since the latter only accounts for about 80 % of all contacts observed in the complexes (see also below) (3, 18). Please note that the K_D_ for the binding of pUL50 to pUL53 is about 6 to 10-fold lower than reported previously by others (31, 33).

Although the binding affinities are comparable across subfamilies, the thermodynamic profiles differ between the three α-, β- and γ-herpesvirus members (Table 1). While in VZV and EBV, the core NEC complex formation is enthalpy driven, complex formation in HCMV is favored by both the enthalpy and entropy terms. The latter might be a consequence of the fact that pUL53 forms homodimers by itself before yielding a heterodimeric pUL50-pUL53. This reduces the loss of molecular degrees of freedom in comparison to complex formation involving Orf27 and BFLF2, which are monomeric in absence of binding partners. Thus, the entropy loss during complex formation can be expected to be smaller in HCMV in comparison to EBV and VZV. The question, whether differences observed in the stoichiometry and thermodynamic parameters in the three prototypical NECs of the α-, β- and γ-herpesviruses investigated here extend to all members of the respective subfamilies, will require further investigations.

### Some degree of binding promiscuity regarding core NEC complex formation within subfamilies

In order to address the question whether the core interaction of the VZV NEC Orf24-Orf27 is highly selective or displays some binding promiscuity with related herpesviral NEC homologs, a CoIP-based interaction analysis was performed. To this end, tagged versions of VZV and other herpesviral NEC proteins were analyzed by transient transfection of 293T cells in pairwise coexpression settings with autologous or nonautologous interaction partners (Fig. 3). This study, in part provides novel data (particularly for the interaction profiles of the VZV NEC proteins Orf24 and Orf27) and in other parts confirms our earlier investigations (regarding several comparative interaction profiles so far collected for α-, β- and γ-herpesviral NEC proteins, (3,18,22)). In the present study, three herpesviral NECs were used for comparison, i.e. α-herpesvirus HSV-1 (pUL34-pUL31), β-herpesvirus HCMV (pUL50-pUL53) and γ-herpesvirus EBV (BFRF1-BFLF2). The CoIP analysis showed a clearly detectable interaction for the VZV-specific pair Orf24-Orf27 (Fig. 3, lanes 3 and 7) as well as the combinations between the two α-herpesviral NEC proteins, i.e. VZV and HSV-1 (lanes 4 and 8). In contrast, the combinations between VZV and β- or γ-herpesviral NEC proteins were negative (lanes 5, 6, 9 and 10). This novel CoIP finding completes our previous investigations and indicates that the capacity of VZV NEC proteins to undergo heterodimeric interactions is restricted within proteins of the α-herpesviral subfamily.

**Figure 3.**
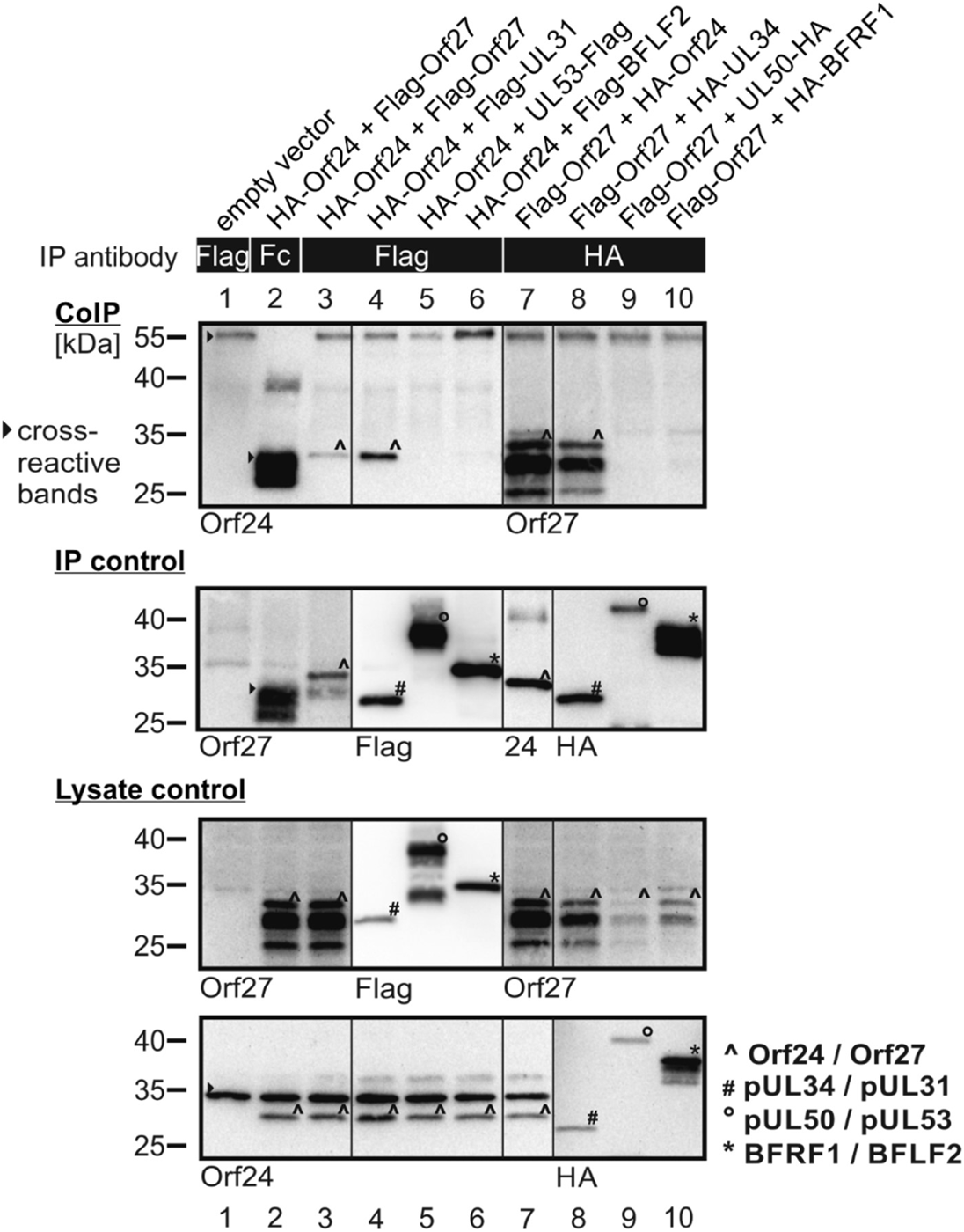
CoIP-based interaction analysis of VZV NEC proteins in pairwise coexpression with autologous or nonautologous herpesviral NEC protein homologs (the latter derived from HSV-1, HCMV and EBV). 293T cells were transiently transfected with expression plasmids coding for HA-tagged and Flag-tagged versions of NEC proteins as indicated. At three d post-transfection (d p.t.), cells were lysed and HA- or Flag-tagged proteins were immunoprecipitated using mAb-Flag, mAb-HA, or a nonreactive antibody Fc fragment as a specificity control. Total lysate controls taken prior to the IP and CoIP samples were subjected to standard Wb analysis using tag-specific antibodies as indicated (lysate control). Successful immunoprecipitation was monitored by Wb staining with the respective tag- or protein-specific antibodies (IP control). Positive CoIP reactions were marked by symbols (˄, #, ჿ, *; see explanation at the lower right) as referring to the identical symbols on the control panels (filled triangle, cross-reactive, non-specific bands, additionally serving as a loading marker). Note the positive signals obtained for CoIP reactions with autologous or nonautologous NEC pairs derived from the same viral subfamily, but negative results obtained for combinations between different subfamilies.

A second approach was used to substantiate this result on the level of individual cells by using confocal imaging of indirect NEC immunofluorescence stainings. To this end, HeLa cells were transfected with similar combinations of tagged versions of the selected α-, β- and γ-herpesviral NEC protein pairs (Fig. 4). As a new approach, extending beyond earlier investigations, the recruitment of the VZV hook protein Orf27 (Fig. 4*A*, panel 4) to the inner nuclear membrane-associated groove protein Orf24 (panel 10) was used as the basis of interaction analysis (3,18,22). Notably, VZV Orf24 (as all herpesviral NEC groove proteins) is always found in a distinct rim-like nuclear envelope localization, while Orf27 (and other herpesviral hook proteins) is only seen in this location after Orf24-Orf27 interaction and rim recruitment. Here, an expected and pronounced nuclear rim colocalization was observed for VZV Orf27 together with VZV Orf24 (Fig. 4*B*, panels 5-8), and likewise HSV-1 pUL31 was recruited by VZV Orf24 to this rim colocalization (panels 9-12). The converse setting, using VZV Orf27 combined with HSV-1 pUL34 (panels 29-32), provided an almost identical rim staining pattern. However, all other combinations between VZV Orf24 or Orf27 with β- or γ-herpesviral NEC protein counterparts lacked a comparable rim staining pattern. As a novel extension of our NEC studies, this finding was now also assessed by a quantitative evaluation of these microscopic samples. This quantitation further substantiated the perfect rim recruitment between the two VZV proteins and a very strong capacity of both, VZV groove protein Orf24 and hook protein Orf27, to undergo interaction with the HSV-1 homologs, but an extremely low tendency of nonautologous, crossviral interactions outside the α-herpesviral subfamily (Table S2). In addition to earlier studies, this statement is supported by an extended comparative analysis of confocal NEC rim recruitment patterns using different heterodimeric combinations with proteins from six herpesviruses, i.e. VZV, HSV-1, HCMV, MCMV, EBV and KSHV (Table S2). Also in this setting, the principal pattern was obtained, in that all autologous combinations with NEC pairs derived from one virus, showed strongly specific and quantitatively high levels of rim interaction patterns (Table S2, upper part, ≥ 95 % of the entire number of signal-positive cells), with very low levels of diffuse localization signals (< 2.5 %). Importantly, also here the potency of herpesviral NEC proteins to undergo nonautologous interactions with viral NEC counterparts within one subfamily was further substantiated (Table S2, middle part, combinations between HSV-1 and VZV; HCMV and MCMV, EBV and KSHV). These nonautologous interaction patterns ranged between 9 ± 2.6 % and 55 ± 2.2 % (perfect nuclear rim colocalization) or < 2.5 % and 32 ± 5.6 % (partial colocalization), respectively. Notably, no nonautologous interaction between NEC proteins of viruses from different subfamilies was detected in any case (Table S2, lower part, < 2.5 %). Combined, these data confirm that the specificity profile of core NEC formation includes a subfamily-restricted capacity of mutual binding.

**Figure 4.**
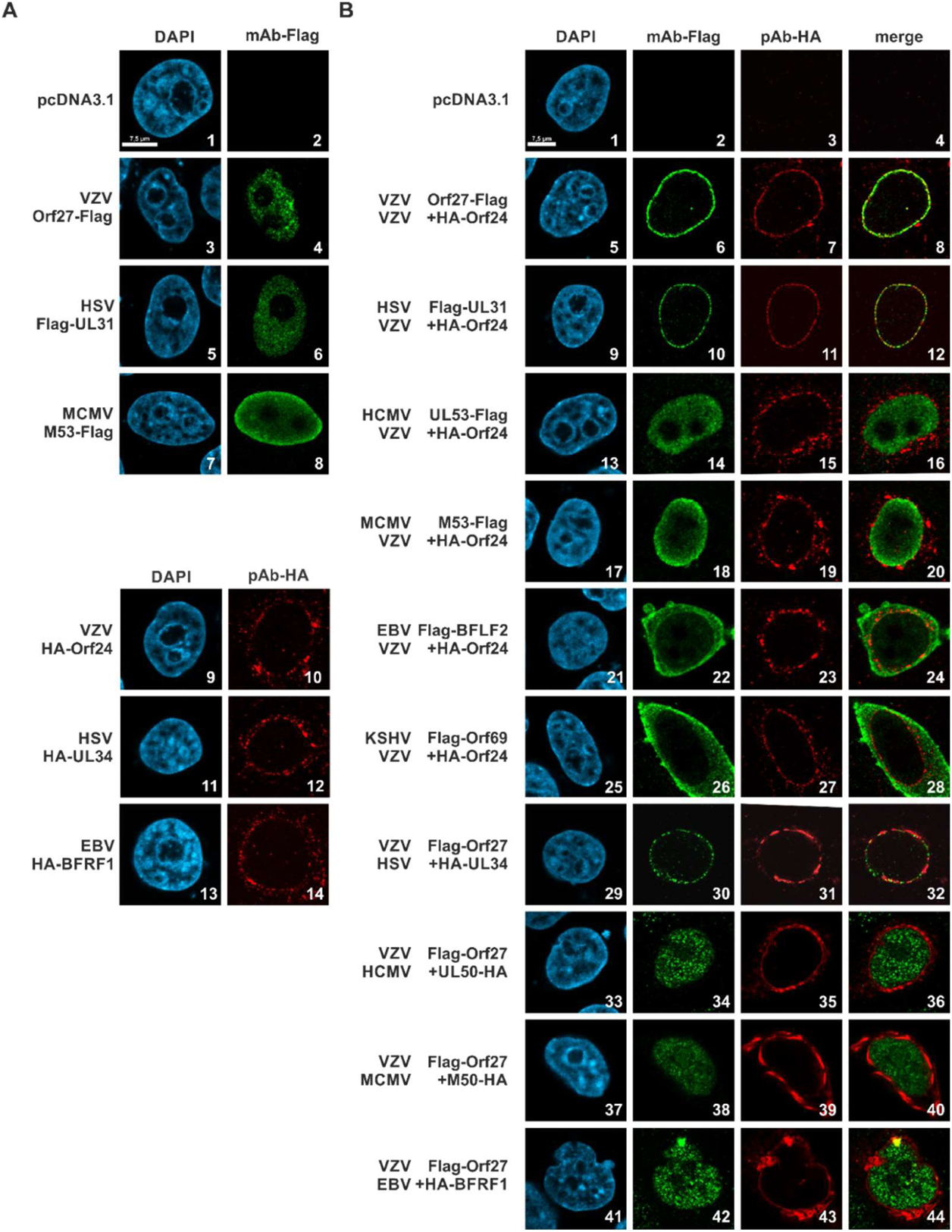
Pairwise transient coexpression of VZV and other herpesviral core NEC proteins. HeLa cells were transiently cotransfected with constructs coding for the tagged versions of NEC proteins. At two d p.t., cells were fixed, used for an immunostaining with tag-specific antibodies and analyzed by confocal imaging. DAPI counterstaining indicated the morphology of nuclei of the respective cells. A, controls of single expression of the VZV NEC hook protein and homologs (panels 1-8) or the VZV NEC groove protein and homologs (panels 9-14). B, pairwise coexpression of the VZV groove (panels 5-28) or hook (panels 29-44) NEC proteins combined with their homologous herpesviral counterparts. Note the perfect nuclear rim colocalization between the VZV NEC proteins alone (Panels 5-8) or NEC counterparts within the same viral subfamily (HSV-1, panels 9-12 and 29-32), which is missing in the combinations with NEC proteins from other subfamilies (HCMV, MCMV, EBV or KSHV)

**Table 2.** Crystallographic data collection, phasing and refinement statistics.

|  |  |
| --- | --- |
| **PDB deposition code** | 7PAB |
| **Data collection** |  |
| Space group | P2<sub>1</sub> |
| Cell dimensions |  |
| *a*, *b*, *c* (Å) | 76.01 35.05 158.22 |
| $\alpha$, $\beta$, $\gamma$ (°) | 90 93.69 90 |
| Wilson B factor (Å<sup>2</sup>) | 51.07 |
| Wavelength (Å) | 0.9763 |
| Resolution (Å) | 19.96 - 2.1 (2.175 - 2.1)<sup>a</sup> |
| *R*<sub>meas</sub> (%) | 14.3 (378) |
| *R*<sub>pim</sub> (%) | 4.1 (105) |
| *I*/$\sigma$(*I*) | 10.2 ( 0.72) |
| *CC*<sub>1/2</sub> (%) | 0.999 (0.463) |
| *CC* * (%) | 1 (0.796) |
| Completeness (%) | 98.97 (99.53) |
| Multiplicity | 11.9 (12.9) |
| **Refinement** |  |
| Resolution range (Å) | 19.96 - 2.1 (2.175 - 2.1) |
| No. of unique reflections | 49198 (4890) |
| Reflections used for *R*<sub>free</sub> | 3679 ( 351) |
| *R*<sub>work</sub> (%) | 21.52 (38.48) |
| *R*<sub>free</sub> (%) | 25.45 (39.18) |
| *CC*<sub>work</sub> (%) | 0.962 (0.687) |
| *CC*<sub>free</sub> (%) | 0.939 (0.674) |
| Ramachandran (%) |  |
| favored/outlier | 97.62 / 0.0 |
| Total no. of atoms | 6675 |
| Protein | 6495 |
| Ligands | 12 |
| Solvent | 168 |
| *B*-factors (Å<sup>2</sup>) |  |
| Average | 79.36 |
| Protein | 79.49 |
| Ligands | 111.93 |
| Solvent | 72.15 |
| No. of TLS groups | 21 |
| R.m.s deviations |  |
| Bond lengths (Å) | 0.002 |
| Bond angles (°) | 0.49 |

<sup>a</sup> Values in parentheses refer to the highest resolution shell.

### Crystal structure of the VZV Orf24-Orf27 complex

The structure of VZV Orf24 in complex with Orf27 was solved at 2.1 Å resolution by crystallizing an Orf24::Orf27 fusion protein in which the C-terminus of Orf24 was fused to the N-terminus of Orf27 *via* a GGSGSGGS linker. This strategy has already been applied to two other NEC complexes before (18). The structure was solved *via* the molecular replacement technique with the HSV-1 core NEC as a search model (PDB entry 4ZXS, (16)) and refined to *R*_work_ = 21.5 % and *R*_free_ = 25.5 % (Table 2, Fig. 5). In the crystals, the asymmetric unit contains two Orf24-Orf27 heterodimers. All bioinformatics analyses were performed from here-on with the A-B and not the C-D heterodimer, since molecule A is considerably better defined than the equivalent molecule C as a result of additional crystal packing contacts and as highlighted by lower B-factors (Fig. S4).

**Figure 5.**
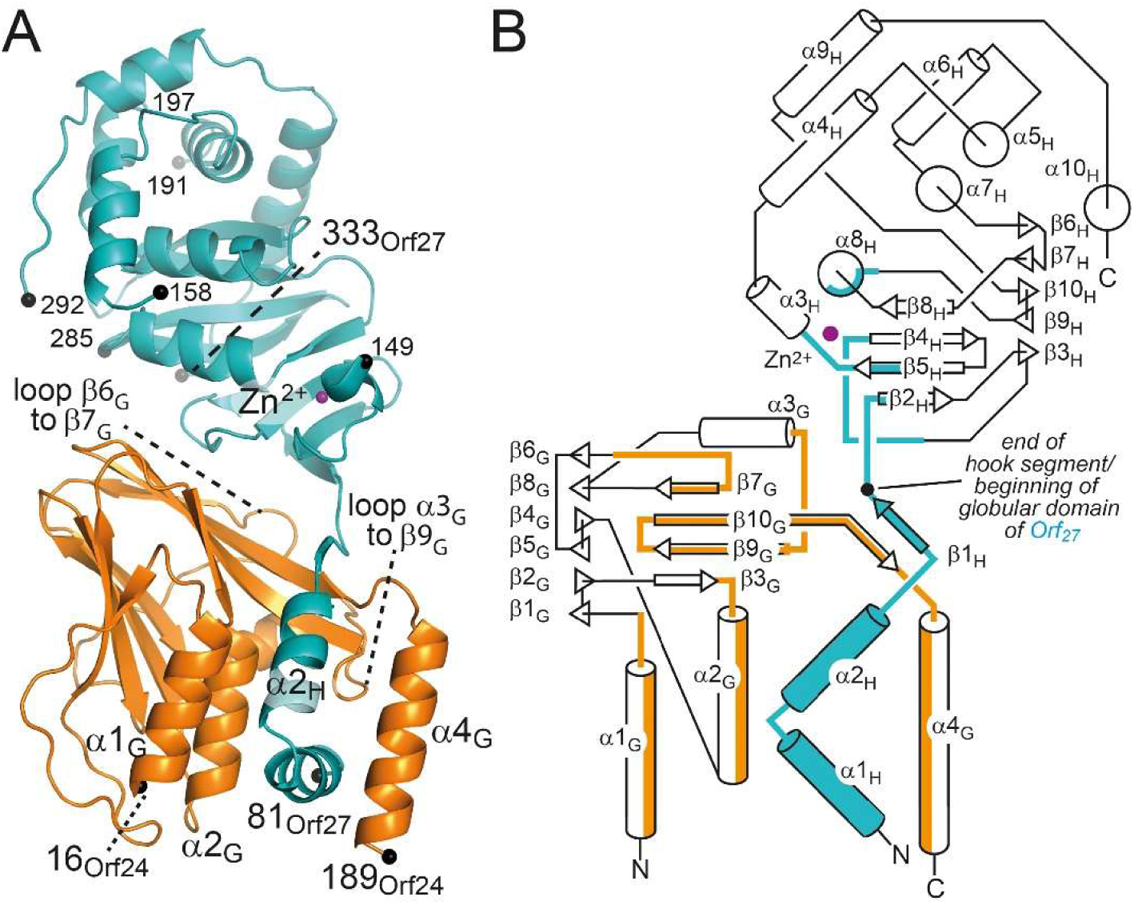
Structure of the VZV Orf24-Orf27 complex. A, ribbon representation of the Orf24-Orf27 complex with Orf24, the groove protein, colored in orange and Orf27, the hook protein, colored in cyan. B, topology diagram of the complex. In B, protein segments involved in Orf24-Orf27 intermolecular contacts are colored. See Table S4 for secondary structure assignments.

The structures of Orf24 and Orf27 closely resemble those of other herpesvirus core NEC proteins present in the protein databank as shown by a DALI webserver search with the individual proteins (Table S3) (34, 35). Orf24 and Orf27 display the highest Z-scores as well as the lowest RMSD_Cα_ values of 1.1 to 1.2 Å when compared to the homologous proteins pUL34 and pUL31 from the α-herpesvirus members PRV and HSV-1. Sequence identities among these α-herpesviral proteins are in excess of 50 % (Table S3). In comparison, RMSD_Cα_ values between 1.9 and 2.8 Å, alongside with sequence identities lower than 21 % are observed when comparing Orf24 and Orf27 to the corresponding proteins from β-herpesviruses, i.e. HCMV and MCMV, or to γ-herpesviruses, i.e. EBV. Obviously, the three α-herpesvirus subfamily members share among themselves a higher sequence and structure similarity than to members from other subfamilies. Orf27 contains a C3H zinc-binding motif with a zinc ion bound to residues Cys128, Cys144, Cys147 and His251. This motif and the overall architecture of the complexes are highly conserved across all NECs (Fig. S5) (19).

### The Orf24-Orf27 interface

As for other NECs, the Orf24-Orf27 interface can be subdivided into two areas, namely the area that involves the hook segment of Orf27 (residues 81 to 109) and the contacts that are formed between the remaining globular part of Orf27 (residues 110 to 333) and Orf24 (Fig. 5). The total contact area amounts to 1825 Å^2^, and of these, 1400 Å^2^ (77 %) are contributed by residues from the hook segment. Similar values have been reported for other α-herpesvirus NECs, namely from HSV-1 and PRV (16, 20). These values also compare well with those observed for the β-herpesvirus HCMV pUL50 and pUL53 complex (1880 Å^2^ in total with 1510 Å^2^ (80 %) contributed by the pUL53 hook segment) (19). In case of γ-herpesvirus EBV, only a structure of BFRF1 in complex with the hook segment of BFLF2 is available, and in this structure, the hook segment contributes 1590 Å^2^ to complex formation (18).

The Orf24-Orf27 complex is formed by interactions involving four segments in each protein (Fig. 6 and Fig. S6). Segment one and two as well as the second half of segment four of Orf24 line the groove into which the hook segment of Orf27 binds. The third segment is formed by residues 113 to 119 of Orf24, and this segment is almost solely involved in interactions with the globular part of Orf27 and in particular with the 121 to 127 segment of Orf27 (see below) (Fig. 6). It should be noted that the fourth Orf24 segment is particularly large. It covers residues 137 to 188 and several secondary structure elements.

**Figure 6.**
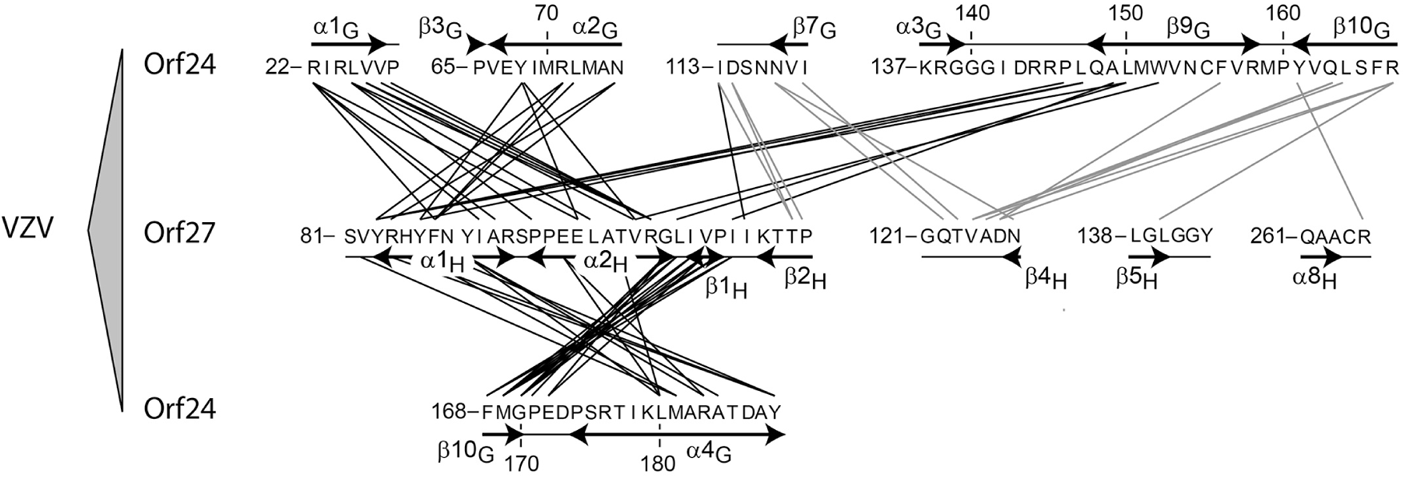
Intermolecular contacts in the VZV Orf24-Orf27 complex. Interacting residues are connected and highlighted by lines based on a 3.8 Å distance cut-off criterion. Interactions involving the hook segment of Orf27 (residues 81-109 containing α1H, α2H and β1H) are marked by black lines and those involving the remaining globular domain of Orf27 by grey lines.

The largest contiguous Orf27 segment participating in the Orf24-Orf27 interface is the segment formed by residues 81 to 113 and which includes the hook segment (residues 81 to 109) (Fig. 6). Remarkably, of the 29 hook residues, all but three participate in binding as judged by the changes in solvent accessible surface areas in these residues upon complex formation. The segment also spans the beginning of β-strand β2_H_ (110 to 113), which contributes an additional two residues to the interface (Fig. 6, Fig. S6 and Table S4). Three additional Orf27 segments also contribute to the interface. Of these, the segment formed by residues 121 to 127 and consisting of the loop β3_H_-to-β4_H_ as well as the beginning of strand β4_H_ appears worth mentioning since also in this segment, all sequential residues participate in the interface. In total, this segment contributes 240 Å^2^ (13.2 % of 1825 Å^2^ total contact area) to the Orf24-Orf27 interface (Fig. S6).

A computational alanine scanning mutagenesis identifies five residues in Orf24 and seven residues in Orf27 that are predicted to contribute more than 2 kcal mol^-1^ to the interaction energy of the complex (Fig. S6). Of these 12 residues, three residues form a continuous interaction patch that is located outside of the hook-into-groove region (Fig. 7*A*). The participating Orf24 residues are Phe156 and Arg167, which are displayed from two different β-strand, namely from strands β9_G_ and β10_G_, respectively. The interacting Orf27 residue is Asp126, which is part of the Orf27 segment 121 to 127 discussed above and which forms a salt bridge with Arg167 from Orf24. Phe156 from Orf24 is located in immediate neighborhood to the salt bridge and forms additional hydrophobic interactions with Orf27 residues Val124 and Ala125. Interestingly, Orf24 residues Phe156 and Arg167 as well as Orf27 residue Asp126 are conserved among α-herpesviruses but are not conserved at homologous positions in β- and γ-herpesviruses (Fig. *S*7).

**Figure 7.**
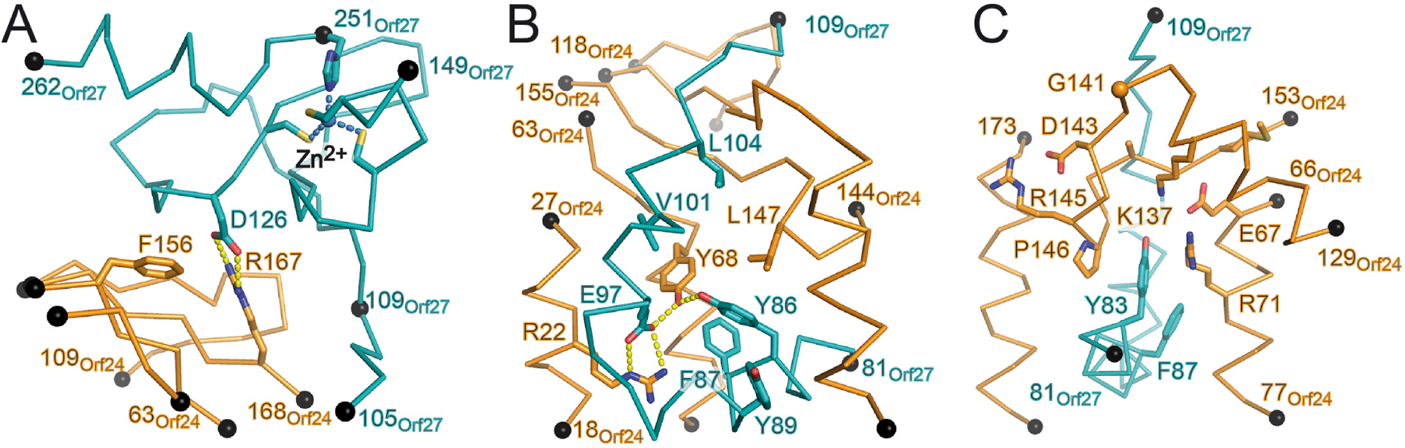
Atomic details of the VZV Orf24-Orf27 interactions. A, interaction patch formed between Orf24 and the globular part of Orf27 as identified by computational Ala-scanning. B, interaction patch formed between Orf24 and the hook segment of Orf27 as identified by computational Ala-scanning. In A and B, interface residues that contribute in excess of 2 kcal mol^-1^ to the interaction are explicitly shown. C, structure of the Tyr83 clamp involving the α3G-to-β9G loop (residues 129 to 153). Residues from this loop that are strictly conserved in a selection of Orf24-homologous proteins are displayed (see also Fig. S7). A selection of additionally conserved residues displayed from other segments of Orf24 are also shown as well as selected Orf27 residues (in cyan). The structural details in A and B are depicted in an orientation that is identical to that of Fig. 5. The orientation in panel C corresponds to that of panel B after a 90° rotation around a vertical axis.

### The hook-into-groove interaction

The hook-into-groove interaction represents the most striking single feature in all available core NECs (3,13,21). Because of the contiguous nature of the hook segment and the concave shape of the complementary binding site in the groove protein, this region shows promise for being targetable by inhibitors. The latter could eventually yield novel antiherpesviral drugs. The side chains displayed by the groove region of Orf24 and the hook segment of Orf27 are chemically highly diverse and display mixed physico-chemical properties (Fig. *S*8). The Orf24 residues that contribute in excess of 75 Å^2^ of their surface to the interaction area are Arg22, Leu25, Tyr68, Met169 and Tyr188. Similarly, the amino acids of the hook segment of Orf27 that contribute more than 75 Å^2^ are Tyr83, His85, Phe87, Tyr89, Val101, Leu104 and Ile108. The intermolecular interactions these residues participate in span from densely packed hydrophobic interactions to extensive networks of polar interactions.

Inspection of the hook-into-groove interface reveals two distinct features, namely (i) an intricate network of polar interactions centered on Orf24 residue Tyr68 and Orf27 residue Tyr86 and (ii) the Orf27 Tyr83 clamp (Fig 7*B* and 7*C*). Computational alanine scanning mutagenesis identified three residues in Orf24 and six in Orf27 that are expected to contribute more than 2 kcal mol^-1^ to the hook-into-groove interaction interface (Fig. *S*6). Interestingly, these residues map almost exclusively to structural feature (i), namely a tyrosine-tyrosine-centered interaction network (Fig. 7*B*). Thus in (i), Orf24 residues Tyr68 (calculated ΔΔG = 5.8 kcal mol^-1^) and Arg22 (4.0 kcal mol^-1^) participate in intermolecular interactions with Orf27 residues Glu97 (4.2 kcal mol^-1^) and Tyr86 (4.0 kcal mol^-1^) through the formation of both a tyrosine-tyrosine interaction and a bidental saltbridge. In addition, Orf27 residues Phe87 (3.1 kcal mol^-1^), Leu104 (2.7 kcal mol^-1^) and Val101 (2.6 kcal mol^-1^) together with Orf24 residue Leu147 (2.3 kcal mol^-1^) extend the interaction patch by adjacent hydrophobic interactions. The only additionally identified residue not participating in this patch is Tyr89 (2.7 kcal mol^-1^); its side chain points in the opposite direction and interacts with helix α4_G_.

In feature (ii), the Tyr83 clamp, the tyrosine side chain displayed from hook helix α1_H_ becomes wedged between Orf24 residues Arg71, Pro146 and Leu147 (Fig. 7*C*). In addition, the hydroxyl group of Tyr83 is in hydrogen-bond distance to the NH-group of Leu147. Not surprisingly, Tyr83 corresponds to the hook residue that contributes most to the interface surface area (135 Å^2^) (Fig. S6*B*). However, its energy contribution to complex formation is estimated to be only 1.5 kcal mol^-1^ (Fig. S6*B*). Of note is that in both features (i) and (ii) many of the participating residues appear to be specifically well conserved in α-herpesviruses and considerably less conserved in β- and γ-herpesviruses (data not shown). Some of the above interactions have been highlighted before in other α-herpesviral NECs (16, 20). However, none of these spotlighted the Tyr83 clamp as a potential determinant for subfamily specificity.

### Structural determinants of subfamily specificity

Although initial analyses revealed many shared features between herpesviral core NEC structures, it has now become increasingly clear that core NEC formation displays subfamily-specific features. While no cross-reactivity has been observed so far between proteins from different subfamilies, nonautologous binding and binding promiscuity can be observed to various extents between NEC proteins from the same herpesviruses subfamilies (Table S2). Analyses of sequence homologies and structure comparisons provide clear hints for the molecular determinants of subfamily-specificity. In an alignment of selected herpesviral NEC groove proteins, which includes all human herpesvirus proteins, three loop regions stand out (Fig. S7). These regions (i) participate in the NEC interface, (ii) differ in length and sequence between subfamilies but at the same time (iii) their composition displays subfamily-specific features.

Loop α3_G_-to-β9_G_ of the groove protein reaches into the space spanned by the two helices forming the hook segment. The loop harbors a salt bridge (between Glu67 and Lys137 in Orf24) that is strictly conserved in all herpesviruses but otherwise differs significantly in sequence and length between subfamilies (Fig. 8*A*). In α-herpesviruses, the loop forms the clamp surrounding Orf27 residue Tyr83. The shortest and longest loops are observed in β- and γ-herpesviral complexes, respectively (Fig. 7*C*, Fig, S7*C*). In both the HCMV and the EBV complexes, the loop sits as a lid on top of α1_H_ rather than embracing its residues as seen with the Tyr83 clamp in α-herpesviruses (Fig. 7*C*, Fig, S9). In all NEC members, the loop α3_G_-to-β9_G_ displays a number of highly conserved subfamily-specific residues (Fig. S7*C*).

**Figure 8.**
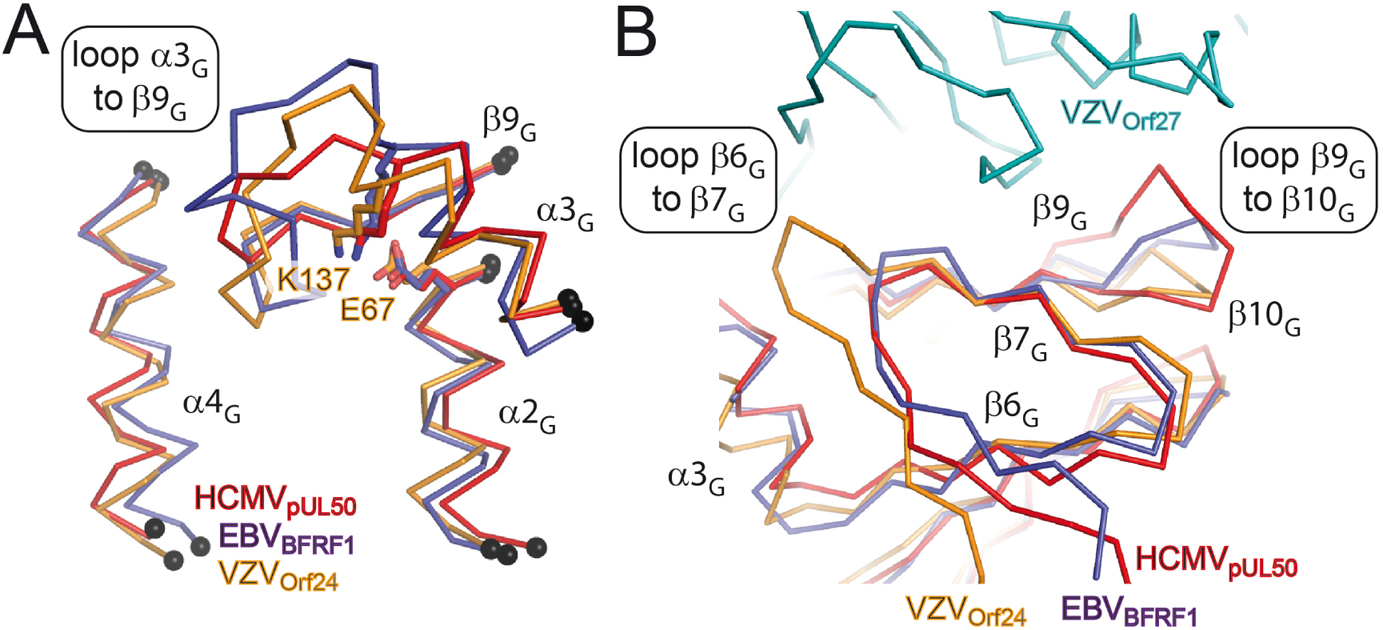
α-, β- and γ-herpesvirus subfamily differences mapped onto three different loop regions. A, differences in the length of the loop α3G-to-β9G appear to be herpesvirus subfamily-specific and give raise to different loop conformations and hook interaction patterns in VZV Orf24 (in orange), HCMV pUL50 (red) and EBV BFRF1 (mauve). The hook segments of the corresponding hook proteins have been omitted for clarity. B, location of two additional loop segments displaying subfamily-specific differences and involved in interactions with the globular domain of the corresponding hook proteins. The orientations of the molecules in all panels correspond to that of Fig. 5 but viewed from the backside.

**Figure 9.**
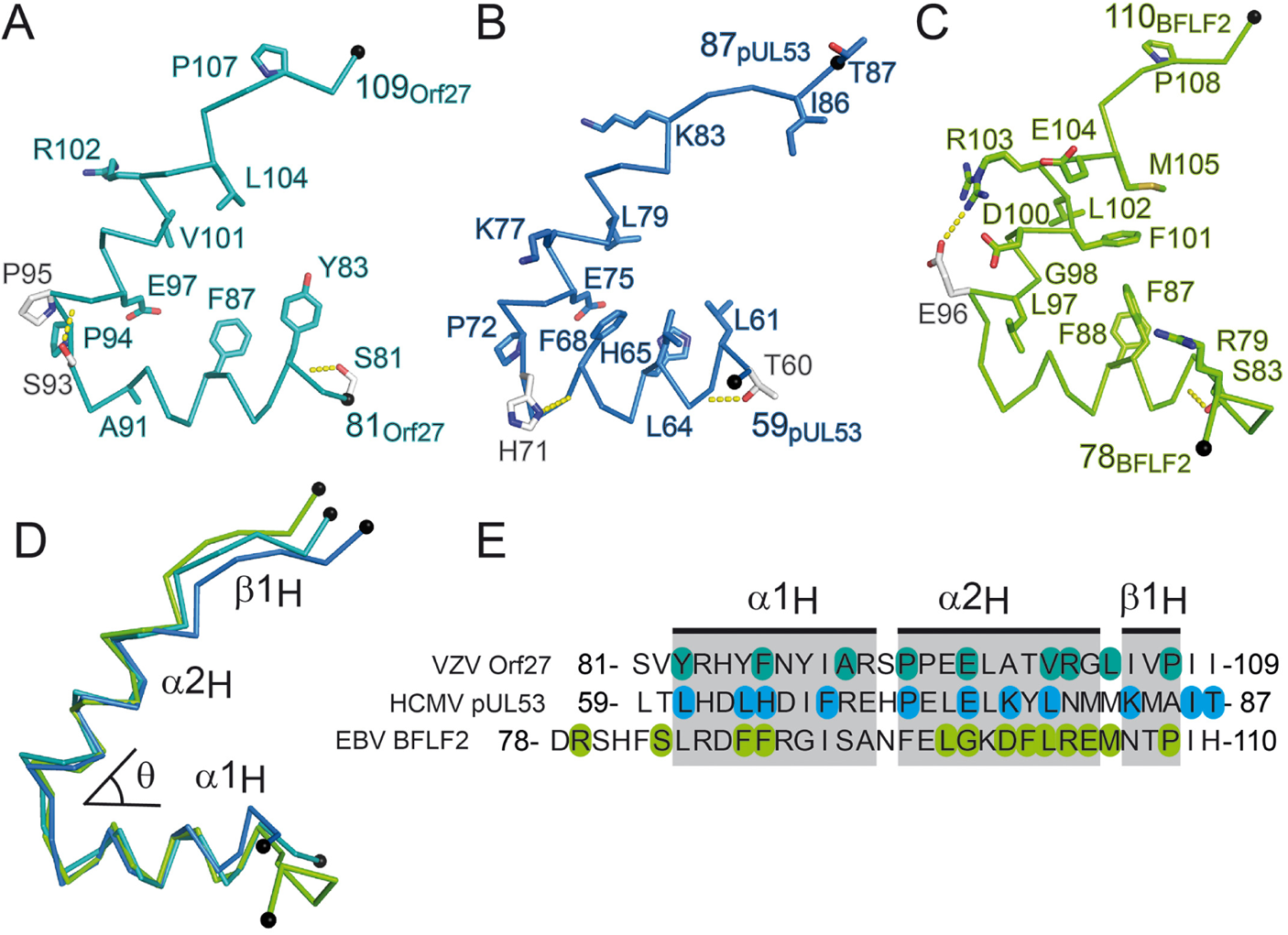
Subfamily-specific differences in the hook segments of VZV, HCMV and EBV. A, hook segment of VZV Orf27 with residues strictly conserved in a selection of α-herpesviral hook proteins displayed in cyan (see also Fig. S7A). B, hook segment of HCMV pUL53 with residues strictly conserved in a selection of β-herpesviral hook proteins displayed in blue. C, hook segment of EBV BFLF2 with residues strictly conserved in a selection of γ-herpesviral hook proteins displayed in green. In A to C, residues displayed in white appear not to be subfamily-specific but contribute interesting features, such as N- or C-terminal helix capping or a surface-exposed salt bridge, to the corresponding hook segment. D, superposition of the hook segments of VZV, HCMV and EBV and definition of the interhelix angle θ. E, sequence alignment of the hook segments. In panel E, the residues explicitly displayed in panels A to C are color-marked. The secondary structure elements are indicated.

Additional subfamily-specific differences can be observed in groove protein loop segments β6_G_-to-β7_G_ and β9_G_-to-β10_G_ (Fig. 8*B,* Fig. S7). The former loop is identical in length in β- and γ-herpesviruses but about five residues longer in α-herpesviruses. In contrast, loop β9_G_-to-β10_G_ is longer by one or two residues in β-herpesviruses in comparison to the other subfamilies (Fig. 8*B,* Fig. S7*D*). In both cases, it appears that the longer loops enable additional contacts with the hook protein that are missing in the subfamilies with the shorter loops. Most likely this will also hold true for γ-herpesviruses albeit so far only a structure of a truncated EBV complex is available.

Subfamily-specific features become also evident when focusing on the hook segments only (Fig. 9). Analysis of the inter-helix angle formed between hook helices α1_H_ and α2_H_ shows that this angle amounts to about 55° and is highly conserved in the α-herpesviral NECs (Table *S*1). An almost identical angle is also observed in the only available γ-herpesviral complex, namely in EBV BFRF1-BFLF2, while an angle of 45° is seen in the β-herpesviral HCMV-specific NEC. Clearly, additional β- and γ-herpesviral NEC structures are needed to corroborate possible subfamily-specific inter-helix angle differences.

In all NECs, the hook structures appear to be stabilized by distinct features. Thus, a structural comparison reveals several instances of helix N-capping such as the N-capping of helix α1_H_ in both Orf27 (involving Ser81) and pUL53 (Thr60) (Fig. 9*A* and *B*). At the same time, His71 in pUL53 forms a C-terminal cap in hook helix α1_H_ in pUL53. While such helix capping residues can be expected to stabilize the conformation of the hook, it is striking that these residues are neither conserved across herpesvirus subfamilies nor within subfamilies (Fig. 9*E*) (36). The hook of BFLF2 from EBV displays also a salt bridge at its surface formed between Glu96 and Arg103 (Fig. 9*C*). However, this salt bridge is neither conserved across herpesviruses nor appears to be subfamily-specific although surface-located salt bridges have been shown to increase the structural stability of proteins (37, 38).

All α- and β-herpesviral hook proteins display a proline residue in the first helical turn of α2_H_ (Fig. 9, Fig. S7*A*). This proline residue might function as a helix breaker thereby disfavoring the formation of a contiguous α-helix spanning both α1_H_ and α2_H_. This could help stabilizing the hook-like conformation of this segment in the prebinding state and hence most likely enhances the binding affinity of the hook. Interestingly, VZV Orf27 even displays two consecutive prolines in this region (Fig. 9*A*). At the same time however, none of the γ-herpesviral hook proteins display a proline at equivalent positions (Fig. S7), and the absence of such a proline residue does not appear to be particularly detrimental to the affinity of EBV BFRF1 to BFLF2 (Table 1).

When comparing the sequences of the hook segment of various herpesvirus hook proteins in detail, then it appears striking that no single residue is conserved across all herpesviruses (Fig 9*E*, Fig. S7*A*). However, a different picture emerges when analyzing the subfamilies individually. In each subfamily, a quite significant number of residues (30 to 40 % of hook residues) can be observed that are strictly conserved within subfamilies including all human herpesviruses (Fig. 9*E,* Fig. S7*A*). When mapped onto the various hook segments, then it becomes obvious that these residues confer the hooks with subfamily-specific surface features. Hence, these features foster subfamily specificity of the hook-into-groove interaction in herpesvirus NECs (Fig. 9).

## DISCUSSION

This report presents novel structural, biochemical and virological data on the core NEC complex of the human pathogenic α-herpesvirus VZV. The main achievements of the study are: (i) a comparative analysis of NEC formation characteristics of three prototypical herpesviruses, namely one from each of the three herpesvirus subfamilies, (ii) the experimental elucidation of the 3D structure of the VZV core NEC Orf24-Orf27 complex, (iii) CoIP and confocal imaging data indicating that Orf24-Orf27 complex formation displays some promiscuity in a herpesvirus subfamily-restricted manner, (iv) computational alanine scanning and structural comparisons highlighting intermolecular interactions shared among α-herpesviruses and (v) an overall structure-function-based model that explains subfamily-specific and common herpesvirus NEC characteristics.

The specificity and selectivity of protein-protein interactions is ruled by protein shape complementarity and the physico-chemical properties of the interacting residues. Any of these may profoundly affect affinities even in the absence of marked changes in the other. In all NEC structures determined so far, the participating proteins display highly similar overall architectures and secondary structure compositions, and complex formation relies on highly similarly shaped binding surfaces. This is particularly well exemplified by the hook-into-groove interaction region, which is built from identical secondary structure elements in all NECs. Yet, the analyses reported here, highlight three loop regions that potentially fine-tune the shape complementarities in a subfamily-specific manner. Of particular interest is the loop α3_G_-to-β9_G_ from the groove protein, which reaches into the interhelical space of the hook (Fig. 7 and 8). Since interaction specificity can arise from both positively and negatively discriminating determinants such as the inclusion of favorable interactions or the preclusion of interactions *via* the introduction of steric clashes, the differences in loop lengths and subfamily-specific sequence identities could allow this loop to function as a shape readout fine-tuning tool and thus represent an important determinant of subfamily specificity.

With the experimental elucidation of a third α-herpesviruses NEC structure, namely that of VZV, it was now possible to analyze with amino acid detail subfamily-specific features and how these α-herpesvirus-specific features differ in the β-herpesviral HCMV NEC complex and in the γ-herpesvirus EBV complex. This reveals a number of individual residue interactions, such as for example the tyrosine-tyrosine interaction network, the Tyr82 clamp and the salt bridge, that are highly conserved in α-herpesviruses and substituted by different sets of subfamily-specific features in β- and γ-herpesviruses. Computational alanine scanning mutagenesis identified twelve highly congruent residues estimated to contribute more than 2 kcal mol^-1^ to the free energy of complex formation. These residues from both the groove and hook protein are in direct contact with each other and participate in two functional hotspots, namely in the formation of the tyrosine-tyrosine interaction network and the salt bridge (Fig. 7). Interestingly, the Tyr82 clamp is not among these hot spots, although Tyr82 contributes as much as 135 Å^2^ surface area to complex formation.

Subfamily-specific patterns of conserved amino acids can also be observed when exclusively focusing on the sequences of the hook segment (Fig. 9). While hook residues that directly interact with respective groove residues appear highly conserved, those residues that help to stabilize the overall conformation of the hook segments, namely helix-capping residues and surface salt bridges are only poorly conserved within and between subfamilies. A partial exception appears to be the presence of a proline at the beginning of α2_H_. Taken together, hook residues that directly participate in groove protein binding appear to be more conserved than residues that merely stabilize the hook conformation.

In light of a broader view on herpesvirus nuclear replication, the core NECs occupy an essential position within viral replication, in that they regulate at least three elaborate processes, namely the multicomponent recruitment of NEC-associated effector proteins, the reorganization of the nuclear lamina and membranes, and the docking of nuclear capsids to the sites of egress. As outlined above, the VZV core NEC displays numerous structural properties closely related to the previously characterized HSV-1 and PRV complexes. As far as the regulatory properties of the VZV NEC are concerned, the question of similar or even identical features in comparison to other α-herpesviral or more distantly related β- and γ-herpesviral complexes remained unclear so far, mostly due to the fact that reports on the VZV NEC and nuclear egress are limited in number to date. So far, little information has been available on whether the associated factors within the VZV-specific multicomponent NEC are shared with other herpesviruses. When comparing the available mass spectrometry-based interactomic data on NEC (for MCMV, see (39); for HCMV, see (40); for HSV-1, see (41)), it appears unlikely that the VZV core NEC binds to identical NEC-associated proteins that are possibly shared within the α-herpesvirus subfamily. This assumption is based on the experimental finding of substantial differences in such binding profiles previously identified for the comparison of the two β-herpesviral representatives HCMV and MCMV. A comprehensive comparison of all so far identified or suggested NEC-associated proteins, both of viral and host cell origin, revealed at least as many matches as differences (13). Thus, it appears very probable that also the VZV-specific NEC may harbour several very distinct, unshared, possibly even strain- or cell type-specific binding partners, which may represent host-directed adaptations and do not stand in contrast to the common theme of NEC functionality found for all herpesviruses (42).

As far as the NEC-facilitated reorganization of the nuclear lamina as well as membranes is concerned, there is no indication to expect a VZV-specific difference. The microscopy-based and protein-protein interaction-based data on VZV-mediated rearrangement of the host cell nuclear envelope published so far are mostly consistent with those provided for HSV-1 (12,15,43–45) or HCMV (13, 46). It should be mentioned, however, that the architecture of neuronal cells preferentially infected by VZV shows specific peculiarities not found by epithelial, lymphoid cells or fibroblasts frequently infected by other herpesviruses, so that also here some host-directed adaptation appears possible. A third point, mentioned with the docking of nuclear capsids to the sites of egress, is very likely to be highly conserved among herpesviruses in a way that the VZV structure described in the present study does not give rise to expected differences. Based on the conserved symmetry shared by α-, β-and γ-herpesviral nuclear capsids, NECs appear to provide a general capsid docking platform at the inner leaflet of the nuclear membrane (13). As a recently accepted model that may hold true for herpesviruses of all three subfamilies, the hexameric arrangement of NEC heterodimers forms a coat-like lattice structure for the docking of nuclear capsids that are also comprising a partially hexameric surface symmetry (13, 47). Although hexameric coat-like NEC arrangements so far have only be detected for α- and β-herpesviruses, i.e. HSV-1, PRV and HCMV, the close structural NEC relatedness among α-herpesviruses described by our present data strongly suggests an identical mode of NEC-capsid interaction also in the case of VZV.

Combined, this report highlights core NEC subfamily specific features in agreement with the observation that core NEC formation displays some promiscuity within subfamilies while no heterologous complexes are formed between subfamilies. In all cases however, complex formation is significantly less efficient in cases heterologous complexes are formed. Because of its central role in herpesviral replication, the inhibition of core NEC formation remains an interesting therapeutic target. The present study suggests that, while it might be possible to identify therapeutics that could possibly inhibit multiple viruses from identical subfamilies, the identification of compounds that are active across subfamily boundaries might be considerably more challenging.

## EXPERIMENTAL PROCEDURES

### Protein production and purification

Plasmids for the expression of HCMV pUL53 residues 50-292, HCMV pUL50 1-175 and EBV BFRF1 1-192 were generated as previously published (18, 19). VZV Orf27 residues 77 to 333 (Uniprot entry: Q6QCM8, (48)) and EBV BFLF2 residues 78 to 318 (Uniprot entry: K9UT32) were each cloned into a pGEX-6P-1 vector (GE Healthcare). The constructs include an N-terminal GST affinity tag and a HRV 3C protease cleavage site (SDLEVLFQGPLGS). All remaining protein constructs were cloned into the pET28b vector (Merck Millipore). Unless otherwise stated, these constructs contain an N-terminal hexa-histidine tag followed by a thrombin cleavage site (MGSSHHHHHHSSGLVPRGSH). For the fusion protein VZV Orf24::Orf27, residues 16 to 189 of VZV Orf24 (Uniprot entry: Q6QCN1) were fused *via* a GGSGSGGS linker to residues 77 to 333 of VZV Orf27. VZV Orf24 by itself consists of residues 16 to 189 with a preceding methionine residue in place of the open reading frame of the pET28b vector. This deletes the original hexa-histidine tag and thrombin cleavage site. Instead, a TEV protease cleavage site and a hexa-histidine tag (ENLYFQGHHHHHH) were fused to the C-terminus of the encoded protein.

Plasmids were transformed into chemically competent *Escherichia coli* BL21(DE3) cells (Novagen) and used to inoculate TB medium supplemented with 100 µg/mL of ampicillin or 50 µg/mL of kanamycin for pGEX-6P-1 or pET28b vectors, respectively. Protein production was induced with 0.25 mM isopropyl 1-thio-β-D-galactopyranoside at 20 °C for approx. 20 h. Cells were harvested and mechanically lysed in appropriate buffers for the subsequent purification steps. VZV Orf24 was purified using a HisTrap (GE Healthcare) affinity chromatography step followed by cleavage with recombinant hexa-histidine tagged TEV protease and a HisTrap recapture step. All other hexa-histidine-tagged proteins were purified using a HisTrap (GE Healthcare) affinity chromatography step followed by thrombin cleavage. Surface lysine residues of VZV Orf24::Orf27 were chemically methylated before crystallization (49). GST fusion proteins were purified using Glutathione Sepharose HP (GE Healthcare) followed by cleavage with recombinant GST-HRV 3C protease and a GST recapture step. If necessary, additional anion exchange chromatography steps were performed. Finally, all protein samples were purified to homogeneity using Superdex 75 prep grade columns (GE Healthcare).

### Complex formation monitored via gel filtration experiments

The *in vitro* complex formation of individually purified NEC proteins was monitored using analytical gel filtration. 200 to 600 µg (in 100 to 500 µL) of equimolar mixtures of the respective NEC proteins from HCMV (pUL50 1-175 and pUL53 50-292), EBV (BFRF1 1-192 and BFLF2 78-318) and VZV (Orf24 16-189 and Orf27 77-333) or 100 to 300 µg samples of individual proteins were injected onto a Superdex 75 10/300 GL column (GE Healthcare) pre-equilibrated with 50 mM TrisHCl, 150 mM NaCl, 1 mM TCEP, pH 8 and eluted with 1.2 column volumes of the aforementioned buffer.

### Isothermal titration calorimetry

For the Isothermal titration calorimetry (ITC) experiments, a Standard Volume Nano ITC (TA Instruments, New Castle, USA) with a 24K gold cell was used. The solutions containing the hook and groove proteins of HCMV (pUL53 and pUL50) and VZV (Orf27 and Orf24) were dialyzed twice against 50 mM HEPES 150 mM NaCl 2.5 mM TCEP pH 7.4, the respective NEC proteins of EBV (BFLF2 and BFRF1) against 25 mM HEPES 150 mM NaCl 2.5 mM TCEP pH 7.4. To minimize the heat generated by buffer mismatch, the hook and groove protein of each pair were dialyzed simultaneously against the same buffer. The groove protein was titrated into the hook protein solution in all experiments (100 µM pUL50 into 14 µM pUL53, 50 µM Orf24 into 8-9 µM Orf27 and 50-90 µM BFRF1 into 8-12 µM BFLF2). Each measurement was performed in triplicates with degassed solutions, consisting of 25 incremental titrations (1 x 5 µL, 24 x 10 µL) interspaced by 480 s time intervals at 25 °C and 150 rpm stirring rate. Blank titrations were performed by titrating each groove protein into buffer. The data were corrected using these and processed with the software NanoAnalyze (v3.11.0, TA Instruments).

### ELISA

High binding Immulon microtiter plates were coated overnight at 4 °C with the hook proteins Orf27, BFLF2 and pUL53, respectively [1 µg/ml (pUL50) and 5 µg/ml (BFLF2 and Orf27), respectively, in 0.1 M sodium carbonate buffer, pH 9.5]. Unspecific binding was blocked with 1% BSA in 0.1 M phosphate buffer, pH 7.2, for 3 h. Plates were then incubated with the His-tagged groove proteins (Orf24, BFRF1 and pUL50, respectively) at two-fold serial dilutions, starting at 586 nM, 531 nM, and 567 nM, respectively, for 3 h. Bound protein was detected using anti-His-HRP conjugate (Sigma; 1:10,000). All proteins and antibodies were in 0.1 M phosphate buffer, pH 7.2, containing 0.1% BSA and 0.01% Tween 20. Plates were washed four times with 0.01% Tween 20 in 0.1 M phosphate buffer, pH 7.2, after each incubation step. Plates were developed with OPD (1 mg/ml) in the presence of 0.03% H_2_O_2_ for approximately 5 min in the dark. After the reaction was stopped with 2 M H_2_SO_4_, absorbance was read at 492 nm.

### Cell culture, virus stocks and infection experiments

Human embryonic kidney epithelial cells HEK 293T and cervix carcinoma epithelial cells HeLa (ATCC) were cultivated at 37 °C, 5 % CO_2_ and 80 % humidity using Dulbecco’s modified Eagle medium (DMEM, 11960044, ThermoFisher Scientific). Primary human foreskin fibroblasts were cultivated in minimal essential medium (MEM), Akata-BX1 cells in RMI 1640 medium (Akata-BX1 EBV-GFP, recombinant expression module selected under 350 µg/ml geneticin supplementation) (50, 51). Cell culture medium was supplemented with 1x GlutaMAX^TM^ (35050038, ThermoFisher Scientific), 10 μg/ml gentamicin and 10 % fetal bovine serum (FBS, F7524, Sigma-Aldrich). Primary human foreskin fibroblasts (HFFs, own repository of primary cell cultures) were propagated as described previously (25, 52) and infectious stocks of HCMV strain AD169-GFP and VZV strain Oka-GFP were produced on HFFs and used for infection experiments according to standard procedures (53–55).

### Transient plasmid transfection

Transient transfection of 293T cells was performed using polyethyleneimine–DNA complexes (Sigma-Aldrich) as described previously (56). HeLa cells were transfected by the use of Lipofectamine 2000 (Thermo Fisher Scientific) according to the manufacturer’s instructions. Plasmids were used as described earlier (18,24–26,29).

### Antibodies

The antibodies used for CoIP, Wb and indirect immunofluorescence (IF) analyses were as follows: mAb-BZLF1 (Santa Cruz, Sc-53904), mAb-IE1p72 (kindly provided by William Britt, University of Alabama, Birmingham, AL, USA), mAb-BFRF1 (kindly provided by Antonella Farina, University of Rome, Italy; (57)), mAb-lamin A/C (ab108595, Abcam, Cambridge, United Kingdom), mAb-HA (Clone 7, H9658, Sigma), pAb-HA (Signalway Eurogentec), mAb-HA-HRP (12013819001, Roche), mAb-Flag (F1804, Sigma), pAb-Flag (F7425, Sigma Aldrich), mAb-Flag-HRP (A8592, Sigma Aldrich), mAb-GFP (11814460001, Roche), mAb-β-Actin (A5441, Sigma Aldrich, St. Louis, MO, USA), anti-mouse Alexa 555 (A-21422, ThermoFisher Scientific), anti-rabbit Alexa 488 (A-11008, ThermoFisher Scientific). Specifically, the series of HCMV-, VZV- and EBV-specific monoclonals, clones: mAb-UL50.01, mAb-UL53.01, mAb-UL97.01, mAb-Orf24 VZ 24.01, mAb-Orf27 VZ 27.01, mAb-Orf47 VZ 47.01, mAb-Orf66 VZ 66.08 and mAb BGLF4.01 was produced by the Center of Proteomics (https://products.capri.com.hr) and the research laboratory of Stipan Jonjic (University of Rijeka, Rijeka, Croatia; (28,40,58,59)). For the generation of mAb-BGLF4.01, recombinant full-length BGLF4 (EBV strain B95-8) was produced and purified from a commercially available Leishmania expression system (Jena Bioscience, Germany; https://www.jenabioscience.com). BALB/c mice were injected subcutaneously with recombinantly expressed protein (50 μg) in complete Freund’s adjuvant. Two weeks later, mice were boosted with the same protein in incomplete Freund’s adjuvant by injecting a two-thirds volume subcutaneously and a one-third volume intraperitoneally. After an additional 2-week period, the sera of immunized mice were screened for antibody titers against the immunogen by using an enzyme-linked immunosorbent assay (ELISA). The best responders were additionally boosted i.p. with the immunogen dissolved in phosphate-buffered saline (PBS). Three days later, spleen cells were collected and, after lysis of red blood cells, fused with SP2/0 myeloma cells at a ratio of 1:1. The cells were seeded onto 96-well tissue culture plates in 20 % RPMI 1640 medium containing hypoxanthine, aminopterin, and thymidine for hybridoma selection. The cultures were screened for mAbs reactive against immunogens by using an ELISA. Positive mother wells were expanded and cloned.

### Coimmunoprecipitation analysis (CoIP)

For CoIP analysis, 293T cells were seeded into 10 cm dishes with a density of 5 x 106 cells and used for transient transfection with expression plasmids. Two to three d post-transfection, CoIP was performed as described previously (26). Antibody-coupled Dynabeads (25 µg/ml, 10002D, ThermoFisher Scientific) were used to obtain specific immunoprecipitates and CoIP samples were further analyzed by standard SDS-PAGE and Western blot (Wb) procedures.

### Indirect immunofluorescence (IF) staining and confocal laser-scanning microscopy

Transiently transfected HeLa cells were grown on coverslips, fixed at 2-3 d p.t. with 10 % formalin solution (10 min, room temperature) and permeabilized by incubation with 0.2 % Triton X-100 solution (15 min, 4 °C). Indirect immunofluorescence staining was performed by incubation with primary antibodies as indicated for 60 min at 37 °C, followed by incubation with dye-conjugated secondary antibodies for 30 min at 37 °C. Cells were mounted with Vectashield Mounting Medium containing DAPI (H-1700, Vector Laboratories, Burlingame, CA, USA) and analyzed using a TCS SP5 confocal laser-scanning microscope (Leica Microsystems, Wetzlar, Germany). Images were processed using the LAS AF software (Leica Microsystems) and Photoshop CS5 (Adobe Inc., San José, CA, USA).

### Crystallization

The fusion protein VZV Orf24::Orf27 was screened for crystallization conditions using the sitting drop technique. Methylated samples were concentrated to 10 and 15 mg/ml in a buffer consisting of 50 mM TrisHCl, 50 mM NaCl, 5 mM DTT and pH 8.0. Crystallization droplets were set up by mixing 2 µl protein and 1 µl reservoir solution. Diffraction quality crystals were obtained at 20 °C with a reservoir solution consisting of 0.09 M Bis-Tris pH 5.5, 0.63 to 0.72 M (NH_4_)_2_SO_4_, 2.7 % PEG 3350 and 4 % formamide. Crystals grew within five to ten days to sizes of 300 x 20 x 10 µm^3^. The crystals were flash-frozen in liquid nitrogen after addition of 20 % ethylene glycol to a protein droplet.

### Data collection and structure refinement

A 2.1 Å resolution diffraction data set was collected at beamline P14 at DESY synchrotron in Hamburg from crystals of VZV Orf24::Orf27. Data were processed with program XDS and datasets from two isomorphous crystals were merged using XSCALE (60). Initial phases were obtained with the molecular replacement technique with program PHENIX_PHASER using the structure of the HSV-1 nuclear egress complex as a search model (PDB entry code 4ZXS) (16, 61). The structure was completed and corrected using either the PHENIX program AUTOBUILD or manually using program COOT (61, 62). Crystallographic data collection and refinement statistics are summarized in Table 2. The model was refined to completion *via* alternating cycles of automated coordinate refinement with PHENIX and manual building with program COOT. The RMSD_Cα_ values were calculated using the DALI webserver (34). All structure illustrations were drawn using Pymol (63). Changes in accessible surface areas were calculated with program AREAIMOL and interhelix angles with program HELIXANG, both from the CCP4 suite (64).

### Computational analyses

The *in silico* alanine scan was done using the PSSM algorithm with default settings of the program FOLD-X (version 5) (65). The multiple sequence alignment was generated with the TEXSHADE package (66). Information from structure-based pairwise alignments was used to ensure a proper alignment in regions of low sequence conservation. For the alignment, the same virus sequences were used as in the previous alignment published in Marschall et al. (3).

## DATA AVAILABILITY

Accession code Protein Data Bank: The coordinates and structure factors have been deposited with the Protein Data Bank under accession code 7PAB.

## ACKNOWLEDGMENTS

The synchrotron MX data were collected at beamline P14 operated by EMBL Hamburg at the PETRA III storage ring (DESY, Hamburg, Germany). We would like to thank Gleb Bourenkov for the assistance in using the beamline. M.M. greatly appreciates the excellent technical assistance by Christina Wangen (Virology, FAU). We are very grateful for our long-term cooperation partners providing very valuable materials and detection tools, i.e. Benedikt Kaufer for the supply with VZV-GFP (Virology, FU Berlin; Ref.), Rona Scott and Lindsey Hutt-Fletcher for providing Akata-BX1/EBV-GFP (Dept. Microbiology and Immunology, Louisiana State Univ. LSUHSC, Shreveport, LA, USA).

## CONFLICT OF INTEREST

The authors declare that they have no conflicts of interest with the contents of this article.

## FOOTNOTES

This research was supported by Deutsche Forschungsgemeinschaft (MA 1289/8-1, EI 423/4-1 and MU 1477/10-1).

## ADDITIONAL INFORMATION

### Supplementary information

***Correspondence*** and requests for materials should be addressed to M.M. and Y.A.M.

## Notes

### Competing Interest Statement

The authors have declared no competing interest.

