## Supplementary material for "The crystal structure of the varicella zoster Orf24-Orf27 nuclear egress complex spotlights multiple determinants of herpesvirus subfamily specificity": VZV_suppl_subm.pdf

**Table S1**

**Compilation of presently available core NEC crystal structures and hook interhelix angles in the respective structures**

| Virus | NEC proteins | PDB entry code (1) | Resol. [Å] | Hook interhelix angle θ [°][a] | |
|---|---|---|---|---|---|
| | | | | Individual observations[b] | Average value |
| α-Herpesviruses | | | | | |
| VZV | Orf24 (16-189)<br>Orf27 (81-333) | present study | 2.1 | 53, 55 | 54 |
| HSV-1 | UL34 (15-185)<br>UL31 (51-306) | 4ZXS (2) | 2.78 | 57, 56 | 56 |
| PRV | UL34 (1-176)<br>UL31 (18-271) | 4Z3U (2) | 2.71 | 55, 55 | 55 |
| PRV | UL34 (4-174)<br>UL31 (26-270) | 5E8C (3) | 2.90 | 55 | 55 |
| β-Herpesvirus | | | | | |
| HCMV | pUL50 (1-175)<br>pUL53 (50-292) | 5D5N (4) | 2.44 | 45 | 45 |
| HCMV | pUL50 (4-168)<br>pUL53 (61-289) | 5DOB (5) | 2.47 | 41 | 41 |
| HCMV | pUL50 (1-171)<br>pUL53 (59-87) | 6T3X (6) | 1.48 | 46, 50 | 48 |
| γ-Herpesvirus | | | | | |
| EBV | BFRF1 (1-192)<br>BFLF2 (78-110) | 6T3Z (6) | 1.56 | 57 | 57 |

[a] Interhelix angle as defined in Fig. 9 and calculated with program helixang from the CCP4 program suite (7).
[b] Angle values reported for the multiple copies present in the crystallographic asymmetric unit of the individual crystal structures (if applicable).

**Table S2**

**Patterns of nuclear rim colocalization of autologous and nonautologous pairs of herpesviral core NEC proteins (VZV marked in grey)[a]**

| Viruses | NEC proteins | Perfect nuclear rim colocalization | Partial colocalization | No colocalization |
|---|---|---|---|---|
| **Autologous combinations** | | | | |
| HSV-1 + HSV-1 | pUL34 + pUL31 | ≥ 95 % | < 2.5 % | < 2.5 % |
| VZV + VZV | Orf24 + Orf27 | ≥ 95 % | < 2.5 % | < 2.5 % |
| HCMV + HCMV | pUL50 + pUL53 | ≥ 95 % | < 2.5 % | < 2.5 % |
| MCMV + MCMV | pM50 + pM53 | ≥ 95 % | < 2.5 % | < 2.5 % |
| EBV + EBV | BFRF1 + BFLF2 | ≥ 95 % | < 2.5 % | < 2.5 % |
| KSHV + KSHV | Orf67 + Orf69 | ≥ 95 % | < 2.5 % | < 2.5 % |
| **Nonautologous combinations within one subfamily** | | | | |
| HSV-1 + VZV | pUL34 + Orf27 | 32 ± 4.7 % | < 2.5 % | 68 ± 4.7 % |
| VZV + HSV-1 | Orf24 + pUL31 | 44 ± 2.5 % | < 2.5 % | 57 ± 2.5 % |
| HCMV + MCMV | pUL50 + pM53 | 53 ± 5.9 % | 21 ± 2.1 % | 26 ± 4.0 % |
| MCMV + HCMV | pM50 + pM53 | 55 ± 2.2 % | 19 ± 1.4 % | 26 ± 2.2 % |
| KSHV + EBV | Orf67 + BFLF2 | 9 ± 2.6 % | 30 ± 2.9 % | 61 ± 5.0 % |
| EBV + KSHV | BFRF1 + Orf69 | 12 ± 3.8 % | 32 ± 5.6 % | 56 ± 9.3 % |
| **Nonautologous combinations between different subfamilies** | | | | |
| HSV-1 + HCMV | pUL34 + pUL53 | < 2.5 % | < 2.5 % | ≥ 95 % |
| HCMV + HSV-1 | pUL50 + pUL34 | < 2.5 % | < 2.5 % | ≥ 95 % |
| HSV-1 + MCMV | pUL34 + pM53 | < 2.5 % | < 2.5 % | ≥ 95 % |
| MCMV + HSV-1 | pM50 + pUL31 | < 2.5 % | < 2.5 % | ≥ 95 % |
| HSV-1 + EBV | pUL34 + BFLF2 | < 2.5 % | < 2.5 % | ≥ 95 % |
| EBV + HSV-1 | BFRF1 + pUL31 | < 2.5 % | < 2.5 % | ≥ 95 % |
| HSV-1 + KSHV | pUL34 + Orf69 | < 2.5 % | < 2.5 % | ≥ 95 % |
| KSHV + HSV-1 | Orf67 + pUL31 | < 2.5 % | < 2.5 % | ≥ 95 % |
| KSHV + HCMV | Orf67 + pUL53 | < 2.5 % | < 2.5 % | ≥ 95 % |
| HCMV + KSHV | pUL50 + Orf69 | < 2.5 % | < 2.5 % | ≥ 95 % |
| KSHV + MCMV | Orf67 + pM53 | < 2.5 % | < 2.5 % | ≥ 95 % |
| MCMV + KSHV | pM50 + Orf69 | < 2.5 % | < 2.5 % | ≥ 95 % |

[a] Comprehensive coexpression and colocalization patterns of a selection of herpesviral core NEC proteins: α-herpesviruses, HSV-1 (pUL34, pUL31) and VZV (Orf24, Orf27); β-herpesviruses, HCMV (pUL50, pUL53) and MCMV (pM50, pM53); γ-herpesviruses, EBV (BFRF1, BFLF2) and KSHV (Orf67, Orf69). HeLa cells were pairwise transiently cotransfected with constructs coding for tagged versions of these proteins. At 2 d p.t., cells were fixed, used for an immunostaining with tag-specific antibodies and analyzed by

confocal imaging. DAPI counterstaining indicated the morphology of nuclei of the respective cells. Areas of at least 50 positive cells were used and all counts were performed in triplicate. The criteria of counting were based on the differentiation of signal-positive cells in those either comprising a perfect nuclear rim colocalization of the two NEC proteins, or a pattern of partial rim colocalization (i.e. including non-rim signals such as dispersed nucleoplasmic staining), or no rim colocalization. Mean values ± SD are given as expressed in a percentage of the entire number of positive cells.

**Table S3**

**VZV Orf24 and Orf27 homologous protein structures as identified using the DALI webserver[a]**

| Rank[b] | Protein name | PDB-entry[c] | Z-score | RMSD (Å)[d] | Nr. of aligned residues/ total nr. of residues present in the deposited model | Identity (%) |
|---|---|---|---|---|---|---|
| Query: Orf24 (16-189, 167 residues)[e] | | | | | | |
| 1 | PRV pUL34 | 4z3u | 27.4 | 1.1 | 169/170 | 57 |
| 2 | HSV-1 pUL34 | 4zxs | 26.1 | 1.2 | 156/159 | 57 |
| 3 | HCMV pUL50 | 5d5n | 16.3 | 2.4 | 149/162 | 19 |
| 4 | EBV BFRF1 | 6t3z | 14.7 | 2.7 | 159/217 | 12 |
| 5 | MCMV M50 | 5a3g | 11.3 | 2.8 | 130/171 | 17 |
| Query: Orf27 (110-333, 205 residues, omitting the hook segment)[f] | | | | | | |
| 1 | HSV-1 pUL31 | 4zxs | 31.3 | 1.1 | 201/241 | 64 |
| 2 | PRV pUL31 | 4z3u | 30.8 | 1.1 | 202/252 | 67 |
| 3 | HCMV pUL53 | 5dob | 24.2 | 1.9 | 192/237 | 21 |

[a] Ref: (8)
[b] In cases where different databank entries described the same protein or contained multiple copies of the same protein, the results are listed for the copy/entry yielding the highest Z-score only
[c] Ref: (1)
[d] Calculated using Cα atom positions.
[e] The query was performed with molecule A of the two copies A and C of Orf24 present in the asymmetric unit of the crystal structure of VZV Orf24::Orf27.
[f] The query was performed with molecule B of the two copies B and D of Orf27 present in the crystal structure of VZV Orf24::Orf27.

**Table S4**

**Secondary structure assignment in the VZV Orf24-Orf27 complex**

| **VZV Orf24 (groove protein, G)** | | **VZV Orf27 (hook protein, H)** | |
|---|---|---|---|
| **Secondary structure element** | **Residues** | **Secondary structure element** | **Residues** |
| $\alpha1_G$ | 16-27 | $\alpha1_H$ | 83-92 |
| $\beta1_G$ | 31-33 | $\alpha2_H$ | 94-103 |
| $\beta2_G$ | 50-57 | $\beta1_H$ | 105-107 |
| $\beta3_G$ | 61-65 | $\beta2_H$ | 110-115 |
| $\alpha2_G$ | 66-77 | $\beta3_H$ | 118-119 |
| $\beta4_G$ | 83-90 | $\beta4_H$ | 127-132 |
| $\beta5_G$ | 93-100 | $\beta5_H$ | 135-140 |
| $\beta6_G$ | 108 | $\alpha3_H$ | 145-149 |
| $\beta7_G$ | 117-121 | $\alpha4_H$ | 160-169 |
| $\beta8_G$ | 124-129 | $\alpha5_H$ | 178-189 |
| $\alpha3_G$ | 130-139 | $\alpha6_H$ | 199-207 |
| $\beta9_G$ | 148-158 | $\alpha7_H$ | 210-220 |
| $\beta10_G$ | 161-170 | $\beta6_H$ | 226-233 |
| $\alpha4_G$ | 174-188 | $\beta7_H$ | 239-245 |
| | | $\beta8_H$ | 249-253 |
| | | $\alpha8_H$ | 254-263 |
| | | $\beta9_H$ | 267-274 |
| | | $\beta10_H$ | 277-285 |
| | | $\alpha9_H$ | 300-308 |
| | | $\alpha10_H$ | 314-326 |

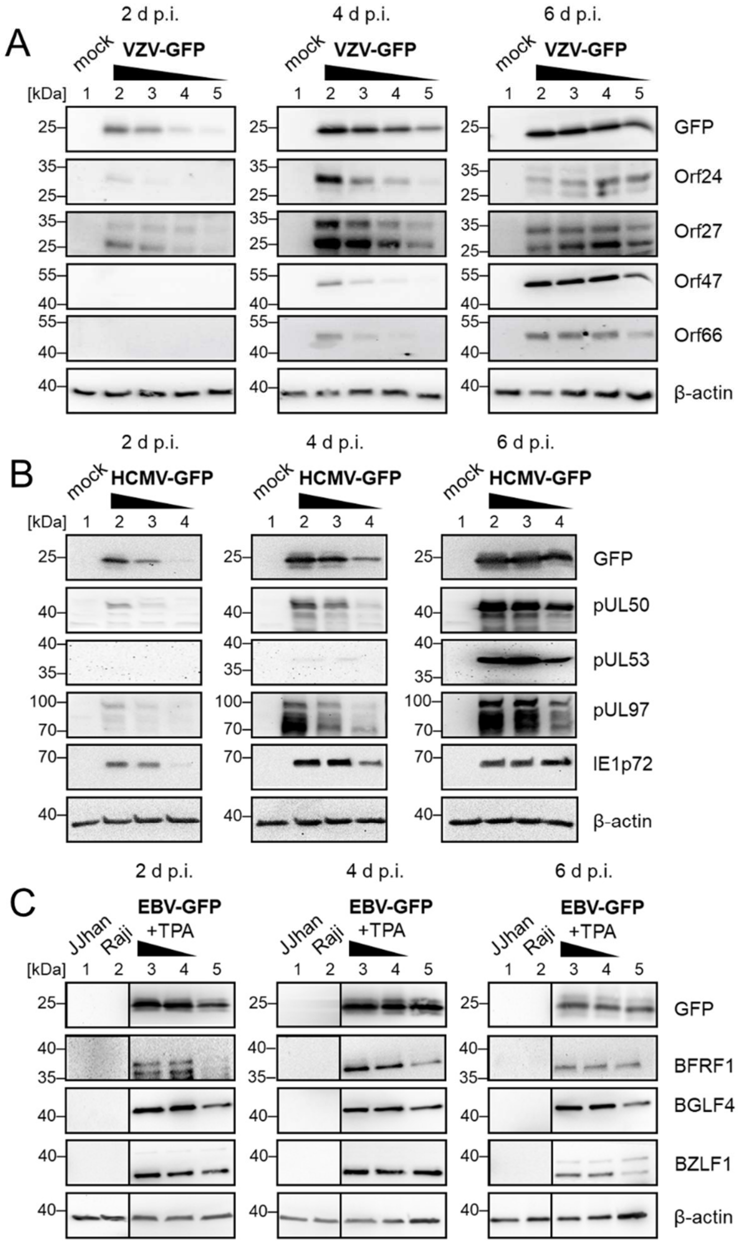


**Figure S1. Core NEC proteins and NEC-associated protein kinases expressed by three human α-, β- and γ-herpesviruses in cell culture infection.**

*A*, The GFP-expressing recombinant of VZV strain Oka (VZV-GFP) was used for infection of primary HFFs by the use of a cell-associated virus inoculum in a range of MOIs 0.5-0.05, or remained uninfected (mock), as described before (9). *B*, The GFP-expressing recombinant of HCMV strain AD169 (HCMV-GFP) was used for infection of primary

HFFs by the use of an infectious supernatant virus inoculum in a range of MOIs 0.5-0.01 (10). *C*, The GFP-expressing recombinant of EBV strain Akata (EBV-GFP) was induced for onset of the viral lytic cycle by chemical stimulation of Akata-BX1 cells with TPA at concentrations of 25, 8 and 0 ng/ml (EBV-positive, latently infected Raji cells and EBV-negative JJhan cells were used as controls lacking the production of lytic viral proteins). Total lysates were prepared at 2, 4 and 6 d p.i. before samples were subjected to SDS-PAGE/Wb separation. Viral proteins and the GFP reporter were immunostained by the specific antibodies as indicated, additional β-actin staining was used as a loading control.

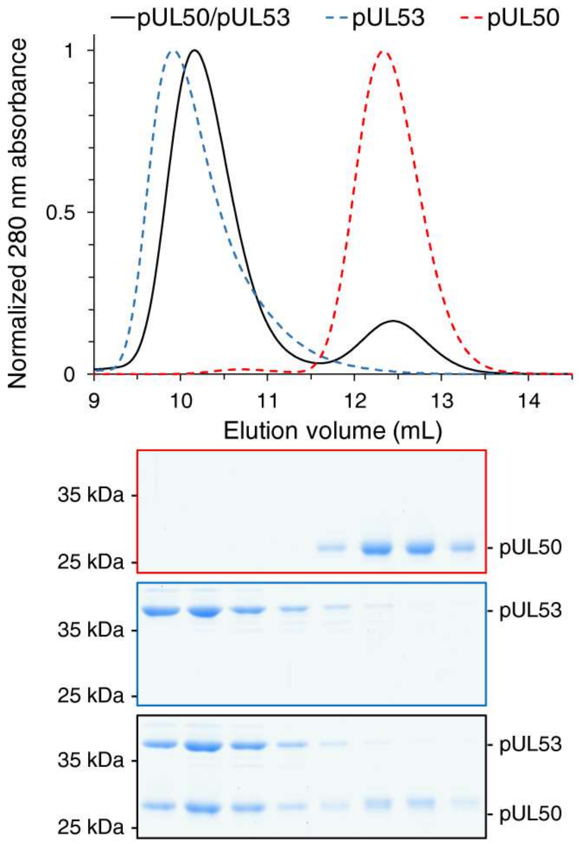


**Figure S2. Stoichiometry of HCMV core NEC formation.**

Core NEC formation in HCMV as analyzed using a gel filtration experiment and SDS-PAGE of peak fractions. *Top panel:* Elution profiles of pUL50 1-175 (red, dashed line), pUL53 50-292 (blue, dashed line) or both (black, solid line). *Bottom panel:* SDS-PAGE of fractions of corresponding elution volumes. pUL50 1-175 (red box), pUL53 50-292 (blue box) or both (black box). These experiments show that homodimeric pUL53 interacts with monomeric pUL50 to form a heterodimeric pUL50-pUL53 complex, as previously reported (11).

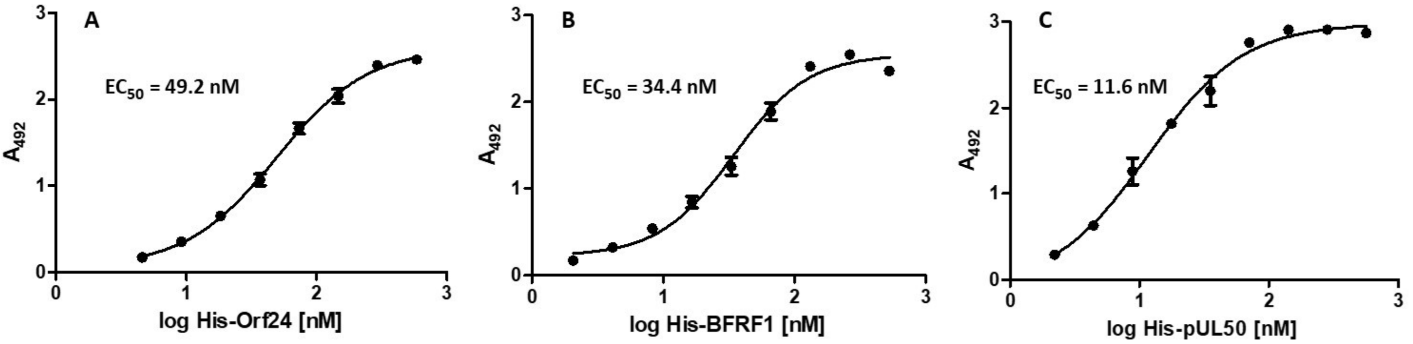


**Figure S3. VZV, EBV and HCMV core NEC interactions assessed by ELISA.**

*A*, VZV Orf24-Orf27 interaction. *B*, EBV BFRF1-BFLF2 interaction. *C*, HCMV pUL50-pUL53 interaction. Immobilized hook proteins were incubated with the respective His-tagged groove proteins at serial dilutions.

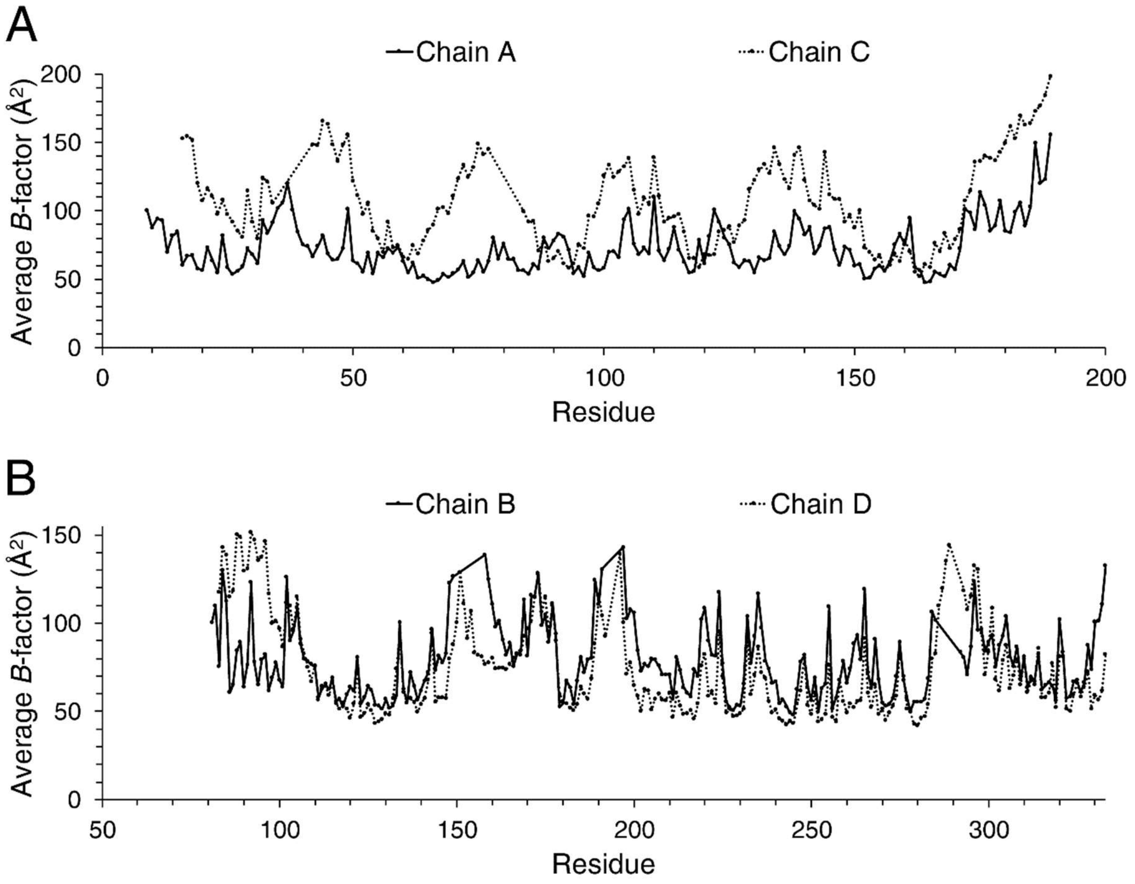


**Figure S4.** ***B*-factor/thermal displacement factor plot of the two copies of the VZV Orf24-Orf27 complex present in the crystallographic asymmetric unit.**

*A*, *B*-factors of the two copies of Orf24 (chain A and chain C). *B*, *B*-factors of the two copies of Orf27 (chain B and chain D). The plot shows that of the two Orf24-Orf27 complexes present in the crystallographic asymmetric unit, the heterodimer A-B is better defined than the heterodimer C-D.

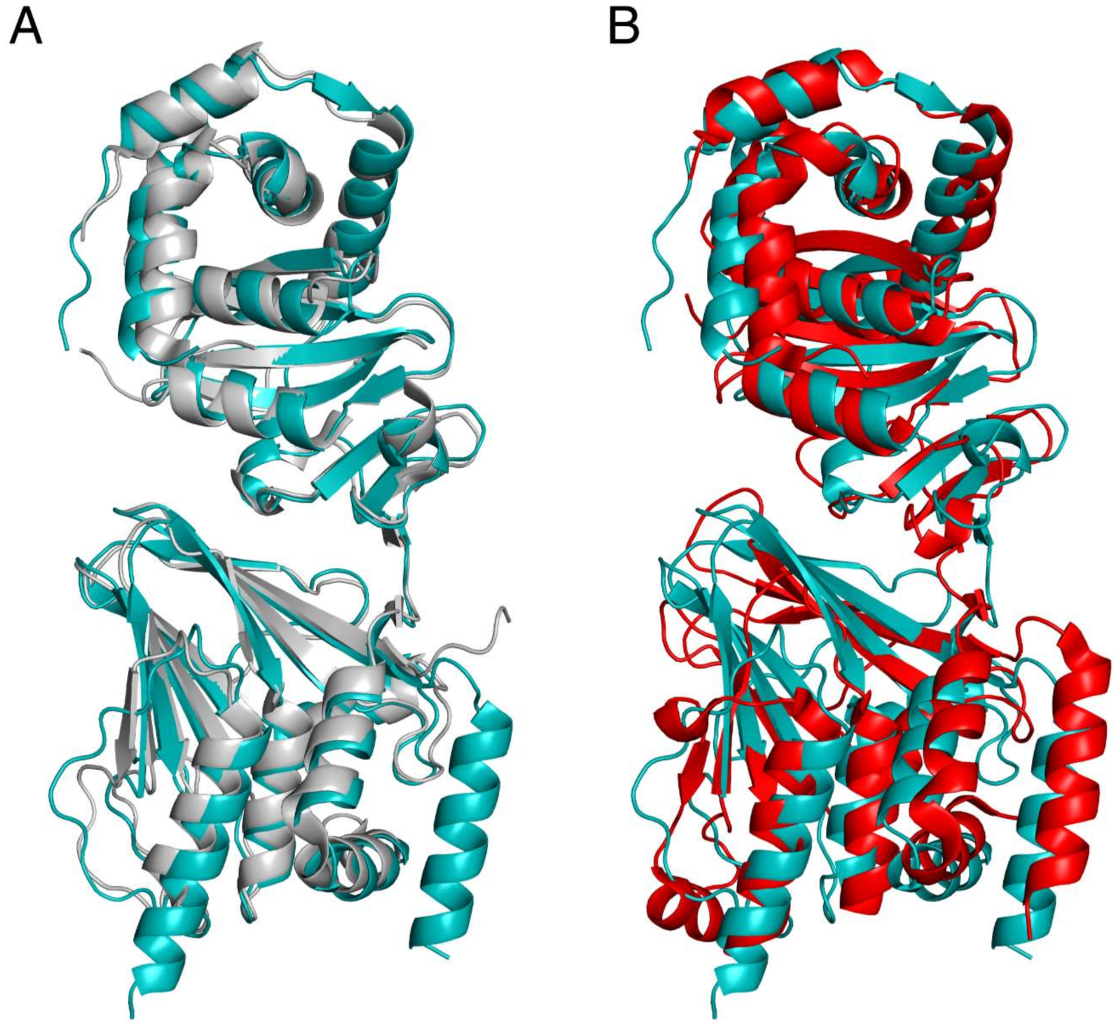


**Figure S5. Superposition of the core NEC structure from VZV with the NEC structure from HSV-1 and HCMV.** *A*, Superposition of VZV Orf24-Orf27 (teal) and HSV-1 pUL34-pUL31 (gray; molecules C and D from PDB entry 4ZXS). *B*, Superposition of VZV Orf24-Orf27 (cyan) and HCMV pUL50-pUL53 (red; PDB entry 5D5N). The structures were superimposed in *Coot* using *SSM Superpose* (12).

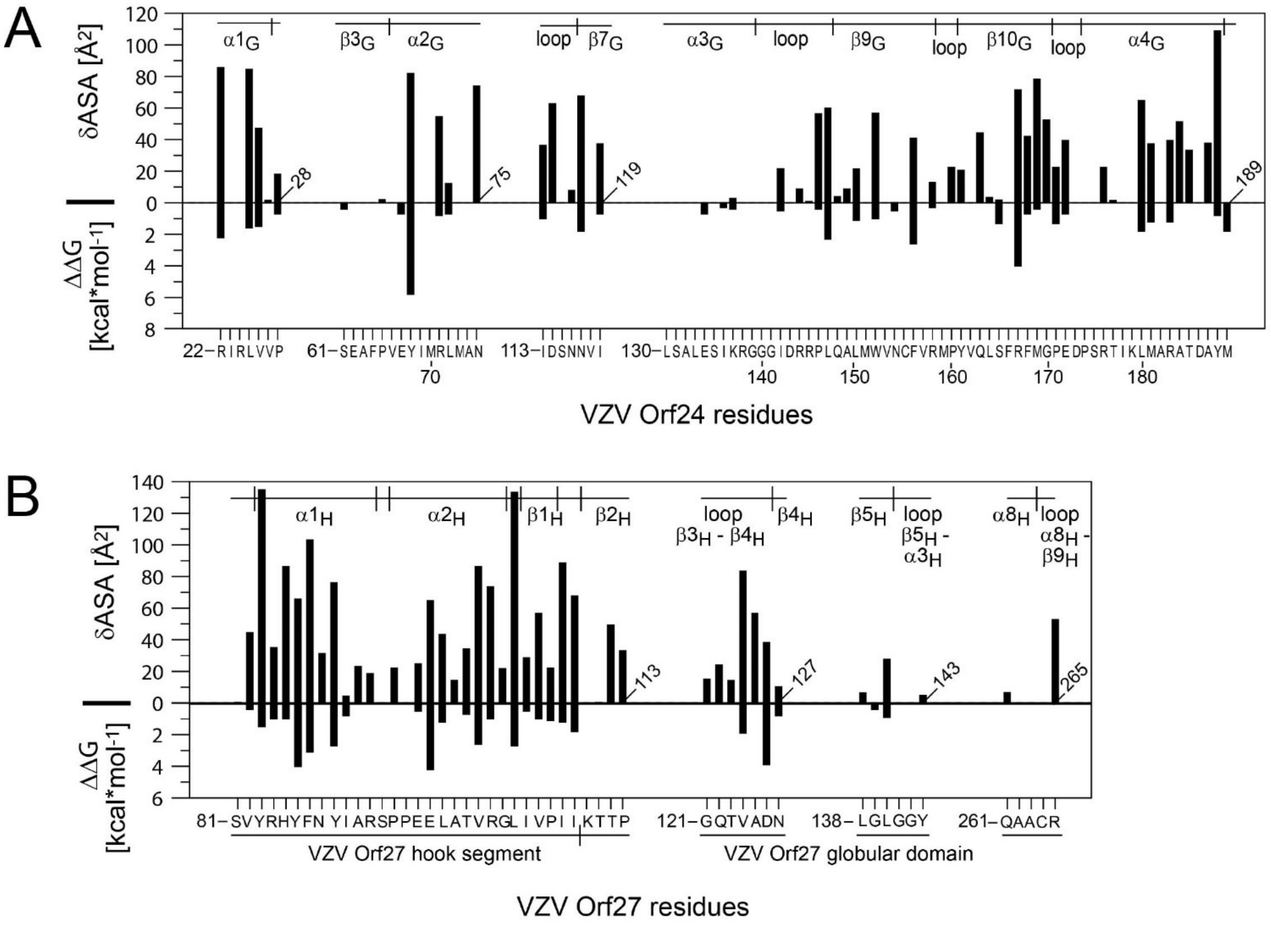


**Figure S6. Surface area and computational alanine scanning contributions of individual residues to Orf24-Orf27 complex formation.**

*A,* changes in the accessible surface area (δASA) and computed ΔΔG values of all Orf24 residues participating in the Orf24-Orf27 interaction. *B*, changes in the accessible surface area and computed ΔΔG values of individual Orf27 residues. Only ΔΔGs larger than 0.2 kcal $mol^{-1}$ are being displayed and discussed (see below). Positive ΔΔG values correspond to a reduction of the interaction energy upon replacing the corresponding amino acid by alanine. Conversely, negative values suggest an affinity improvement. In Orf24, three residues displayed negative ΔΔG values smaller than -0.2 kcal $mol^{-1}$, namely Asn75, Asn116 and Gly170 with values ranging from -0.4 to -0.7 kcal $mol^{-1}$. No such residues were observed in Orf27.

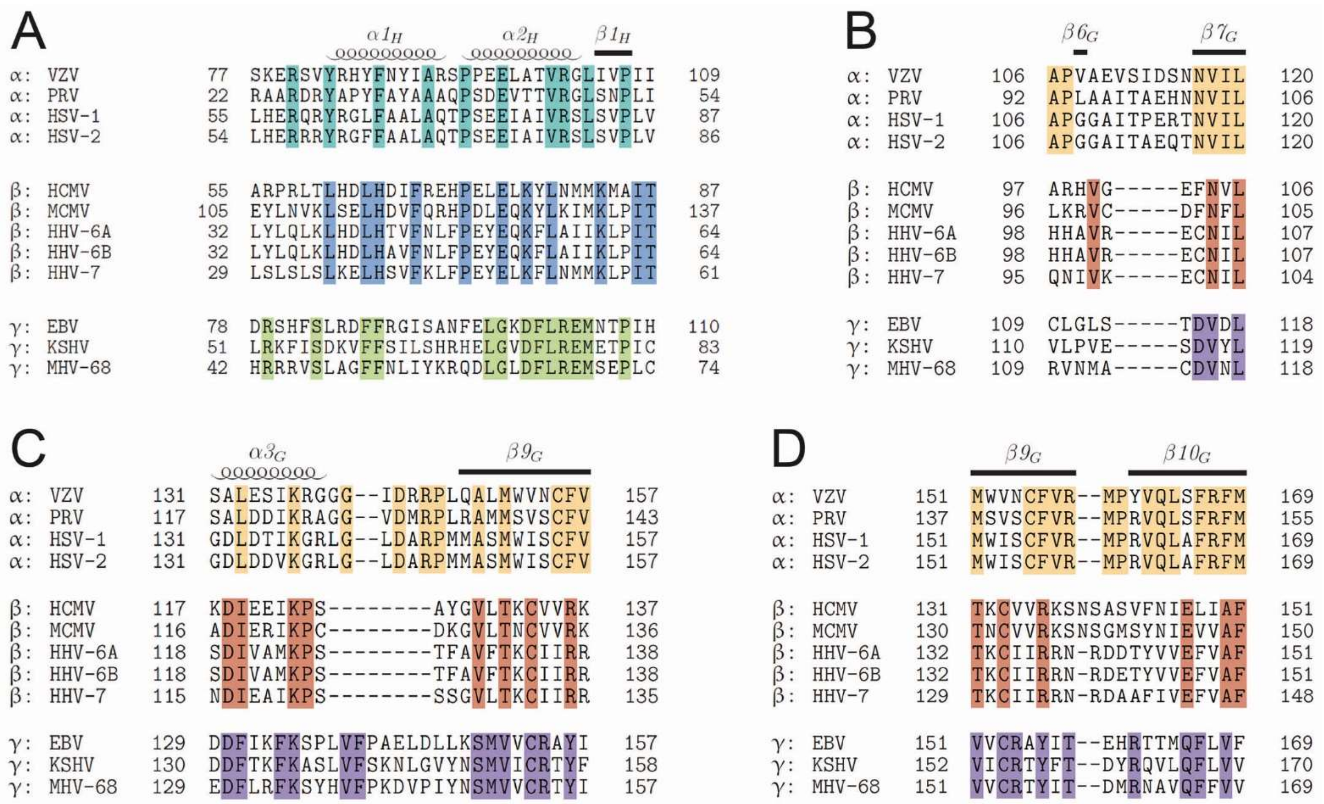

**Figure S7. Sequence conservation in selected segments of α-, β- and γ-herpesvirus hook and groove proteins.**

*A,* structure-based sequence alignment of the hook segment of selected α-, β- and γ-herpesvirus hook proteins (including all human herpesviruses). *B,* structure-based sequence alignment of loop β6$_G$-to-β7$_G$, *C*, loop α3$_G$-to-β9$_G$ and *D*, loop β9$_G$-to-β10$_G$ of selected α-, β- and γ-herpesviral groove proteins. The secondary structure elements from the VZV crystal structure are indicated on top of the alignment. Sequence positions that are strictly conserved within the subfamilies are highlighted by color.

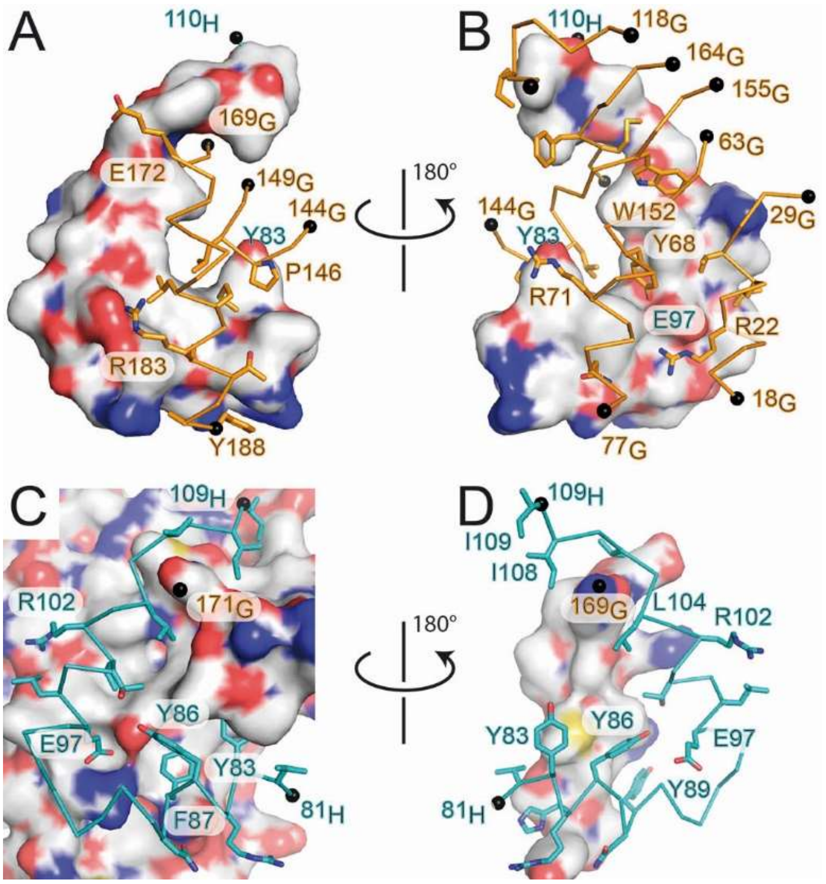


**Figure S8. Display of the hook-into-groove interface.**

*A* and *B*, surface representation of the hook segment of Orf27 in two different orientations with residues from Orf24 displayed in orange and in a stick representation. Only those Orf24 residues are displayed that contribute more than 30 Å$^2$ of their surface area to the interaction interface. *C* and *D*, surface representation of Orf27 viewed from two orientations of the hook segment. The Cα-trace of the hook segment is colored in cyan and hook residues contributing in excess of 30 Å$^2$ of their surface to the interaction interface are displayed (see also Fig. S6).

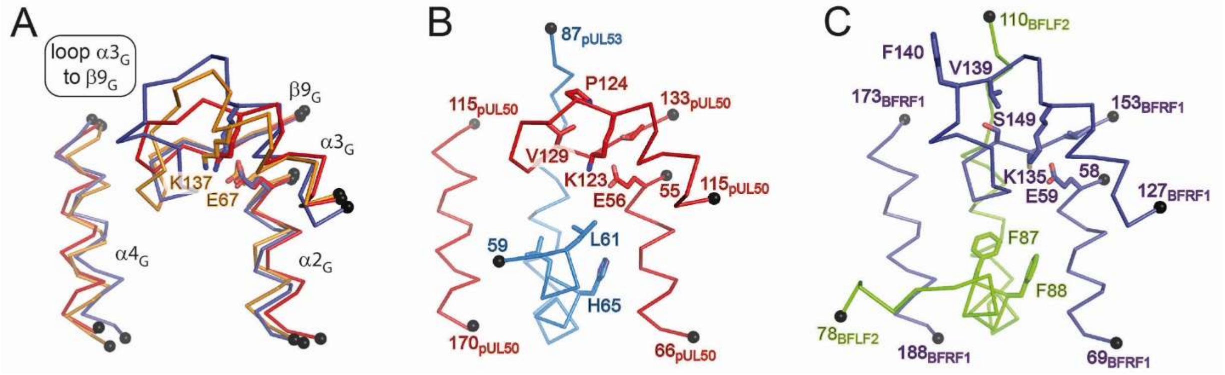


**Figure S9. α-, β- and γ-herpesvirus subfamily differences in loop α3$_G$-to-β9$_G$.**
*A*, superposition of loop α3$_G$-to-β9$_G$ as observed in Orf24 (in orange), pUL50 (red) and BFRF1(mauve). *B*, details of the interaction of loop α3$_G$-to-β9$_G$ in pUL50 with the hook segment of pUL53. Residues from this loop that are strictly conserved in a selection pUL50-homologous β-herpesvirus proteins are displayed together with a selection of conserved residues from other segments (13). *C*, details of the interaction of loop α3$_G$-to-β9$_G$ in BFRF1 with the hook segment of BFLF2. Residues from this loop that are strictly conserved in a selection BFRF1-homologous γ-herpesvirus proteins are displayed together with a selection of conserved residues from other segments (13).
